# Cryo-EM Structure of Duck Secretory IgM Reveals a Conserved Pentameric Assembly with Avian-Specific Features at Molecular Interfaces

**DOI:** 10.64898/2026.08.26.747385

**Authors:** Rebecca M. Schneider, Qianqiao Liu, Beth M. Stadtmueller

## Abstract

IgM is the most ancient antibody isotype, playing an important role in both circulatory and mucosal immune responses across vertebrates, yet structural characterization of its polymeric forms is limited outside of mammals. Here, we report the cryo-electron microscopy structure of mallard duck secretory (S) IgM at 3.37-Å resolution. The structure revealed a pentameric core globally similar to human SIgM, supporting the view that pentameric IgM is subject to strong evolutionary constraints. However, compared to mammalian structures, we observed species-specific differences at molecular interfaces. Surface plasmon resonance binding assays characterizing secretory component (SC)-IgM interactions supported structural observations and, when compared to IgA binding, revealed isotype-specific contributions from the avian SC N-terminal extension. Together, these findings establish a comparative structural framework for polymeric IgM across vertebrates and provide insight into how avian SIgM-specific features may support mucosal immunity in birds.

## INTRODUCTION

Immunoglobulin (Ig) M, the evolutionarily oldest antibody isotype, is expressed by jawed vertebrates and plays critical roles in both circulatory and mucosal immunity^1^. In mammals, IgM is the first antibody produced during a primary immune response^2^ and its predominant form is pentameric, comprising five IgM monomers and one joining chain (JC). Circulating in serum, pentameric IgM provides high-avidity antigen binding and initiates complement fixation through the classical pathway^3^. A subset of pentameric IgM is also transported across epithelial barriers to mucosal surfaces by the polymeric (p) immunoglobulin (Ig) Receptor (pIgR), whose ectodomain, secretory component (SC), remains bound to IgM, forming secretory (S) IgM^4–7^. The role of IgM in bridging innate and adaptive immune responses across multiple tissue compartments is thought to be highly conserved^1,8–10^. This broad functional relevance has likely constrained IgM structure; however, host-pathogen coevolution has also driven species-specific adaptations, and the balance of structural conservation versus divergence across vertebrates remains poorly understood. To date, structural characterization of polymeric IgM is limited outside of mammals. Avian IgM is of particular interest, because avian immune system genes have diverged substantially from mammalian counterparts^11^ and species such as the mallard duck (*Anas platyrhynchos*) serve as natural reservoirs for zoonotic pathogens, including influenza A viruses^11–25^.

In mammals, each IgM monomer comprises two heavy chains (HCs) and two light chains (LCs), with each HC containing one variable domain (V_H_) followed by four constant domains (Cμ1, Cμ2, Cμ3, Cμ4) and an eighteen-residue C-terminal tailpiece (Tp), and each LC containing one Ig variable domain (V_L_) and one Ig constant domain (C_L_). The two HCs and two LCs form two antigen-binding fragments (Fabs) and one fragment crystallizable (Fcμ), the latter of which mediates Fc Receptor (FcR)-dependent effector functions. High-resolution structures of human SIgM have revealed SC asymmetrically bound to a pentameric core comprising five Fcμ subunits and one JC, which adopt hexagonal symmetry with the JC occupying the gap between the first and fifth subunit^26,27^. Subsequent structures of human IgM in complex with the IgM Fc Receptor (FcμR) and the IgM-associated protein CD5L have further defined the molecular interfaces mediating IgM effector functions^28–33^.

IgA is another isotype found in birds and mammals that can be assembled into JC-containing polymeric Igs (pIgs) and transported to mucosal surfaces by pIgR, where it serves as the predominant secretory antibody^34–36^. Cryo-electron microscopy (cryo-EM) structures of mammalian SIgA established conformational asymmetry as a defining feature^37–40^ and subsequent work has extended structural characterization to IgA complexes with host receptors and pathogen-derived factors^41,42^. Additionally, we recently reported the cryo-EM structures of mallard duck IgA and SIgA (*ap* FcαJ and *ap* SFcαJ, respectively), revealing a tetrameric organization distinct from mammalian dimeric IgA, as well as avian-specific features including a four-domain SC (compared to five in mammals) and extended N-termini on SC and JC (i.e. SC_N-ext._ and JC_N-ext._, respectively), which expand the duck SC-IgA interface relative to mammalian SIgA^43^.

To investigate IgM species– and isotype-specific differences and similarities we determined the cryo-EM structure of mallard duck SIgM (*ap* SFcμJ). Comparative analysis with human SIgM and avian SIgA structures, along with surface plasmon resonance (SPR) binding assays, revealed conserved features of the pentameric IgM core alongside species-specific differences at interfaces involving adjacent Fcμ subunits, the JC, and the SC. Together, these findings expand the comparative structural framework for polymeric immunoglobulins across vertebrates and provide insight into how IgM structure has evolved to support conserved and divergent immune functions in birds.

## RESULTS

### The cryo-EM structure of avian SIgM

To investigate the structure of avian SIgM, we co-expressed mallard duck, *Anas platyrhynchos* (*ap*), HC (containing Cμ2-Cμ3-Cμ4-Tp), JC, and pIgR ectodomain (SC; residues 1-458) constructs and determined the cryo-EM structure, *ap* SFcμJ, to an average resolution of 3.37 Å (Fig. 1a, Supplementary Fig. 1, 2a-e, Supplementary Table 1). Local resolution was variable (Supplementary Fig. 2f-j); however, density for most main chain and side chain atoms at molecular interfaces were well-resolved. Local resolution of Fcμ1 and Fcμ5 Cμ4 residues was higher (3.0 Å to 3.7 Å) compared to Fcμ2, Fcμ3, and Fcμ4 Cμ4 residues (3.2 Å to 4.9 Å). Cμ3 domain residues outside of molecular interfaces were less resolved with the local resolution ranging from 3.7 Å to 7.2 Å; main chain and side chain atoms for Cμ3 domain residues with lower resolution were built in geometrically reasonable positions. Consistent with poorly resolved Cμ2 domains in human IgM structures^26–31,44^, *ap* SFcμJ Cμ2 domains were largely disordered and were not built. A subset of HC C-terminal Tps residues (431-448) were also disordered (see below). Local resolution for JC residues varied with interfacing residues ranging from 3.4 Å to 3.6 Å and solvent-exposed residues in loops and the four N-terminal residues ranging from 3.8 Å to 5.7 Å. Local resolution for SC domains (D1-D4) ranged from 3.0 Å to 6.6 Å with the lowest resolution observed in D4.

**Figure 1.**
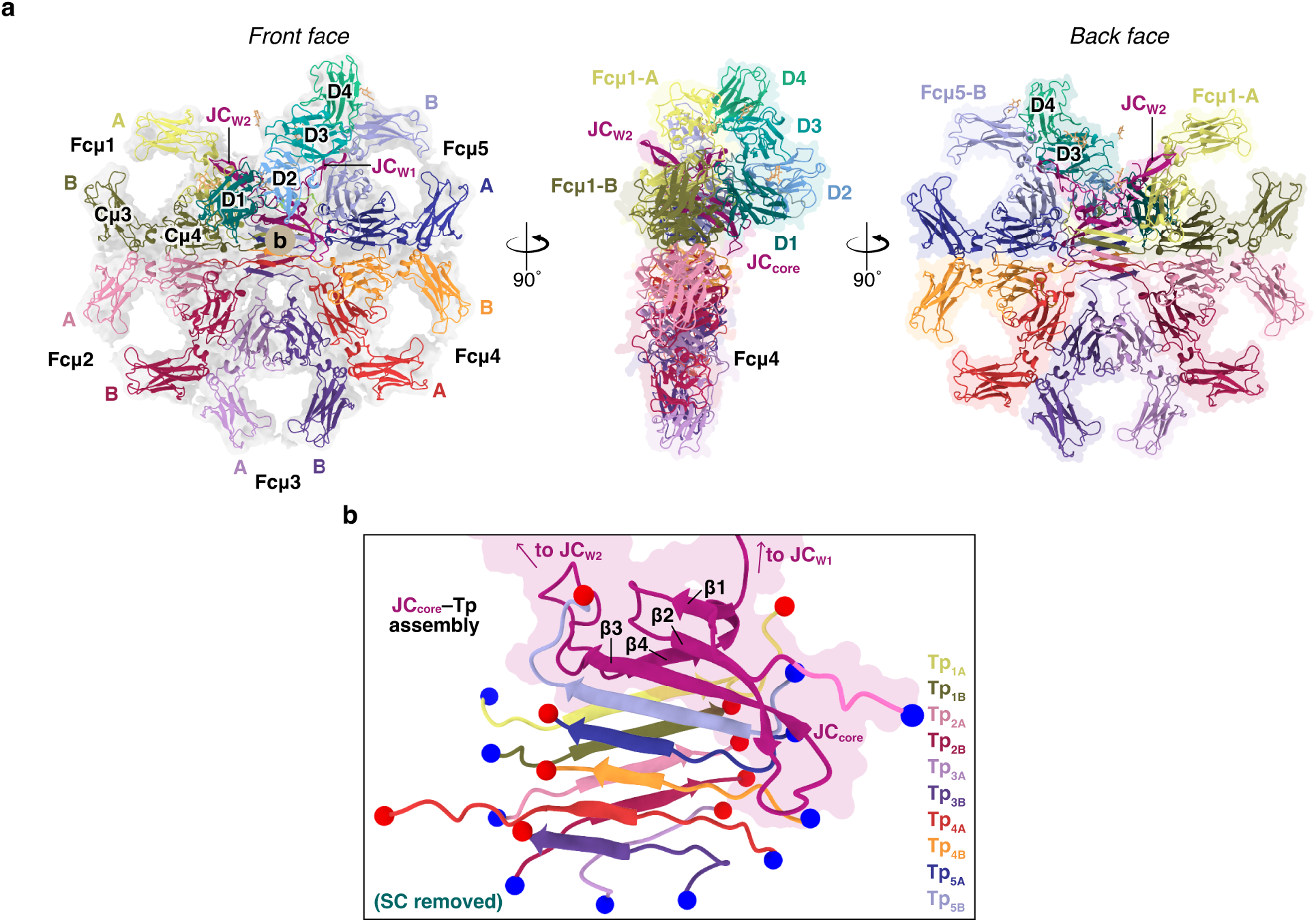
The structure of *ap* SFcμJ. (**a**) Front, side, and back face views of *ap* SFcμJ shown as cartoons with glycans shown as sticks; HCs, Fcμ subunits (Fcμ1-Fcμ5), JC, and SC domains are uniquely colored and labeled. The front face view also shows a transparent density map; side and back face views also show flat surface representations. (**b**) Front face cartoon of JC-Tp assembly with JC_core_ also shown as a flat molecular surface; chains are colored as in (a) and labeled along with the four JC_core_ β-strands. Tp N– and C–termini are indicated by blue and red spheres, respectively.

The refined *ap* SFcμJ structure revealed a pentameric FcμJ core with five Fc subunits (Fcμ1 to Fcμ5, each comprising two HCs designated A and B) and one JC, which was asymmetrically bound by SC. The structure exhibited distinct front and back faces and was globally similar to published human SIgM structures (Fig. 1a). The Fcμ subunits are arranged such that each subunit forms an interface with one to two adjacent subunits, resulting in four Fc-Fc interfaces (i.e. Fcμ1-Fcμ2, Fcμ2-Fcμ3, Fcμ3-Fcμ4, Fcμ4-Fcμ5). At the center of the complex, ten HC tailpieces (Tp_1A_ to Tp_5B_) form a β-sandwich-like domain that is further extended by JC β-strands β1, β2, and β3, which extend the assembly on the front face, and β4, which extends the assembly on the back face (Fig. 1b). These strands, referred to as the JC_core_, also make contacts with residues in Fcμ1 chain A (Fcμ1-A), Fcμ4-B, and Fcμ5-B (Fig. 1a). Two β-hairpins, or ‘wings’^38^, JC_W1_ and JC_W2_, comprise the second half of the JC sequence. Unlike previously reported human SIgM structures^26,27^, *ap* SFcμJ JC_W1_ was ordered and observed contacting Fcμ5-B and SC (Fig. 1a), comparable to that observed in the recently reported avian SFcαJ structure^43^. While JC_W1_ extends from the center of the complex toward the front face, JC_W2_ extends from the center of the complex toward the back face and contacts Fcμ1-A, which also forms an interface with the JC C-terminus (Fig. 1a). The structure also revealed SC asymmetrically bound to the front face of the FcμJ complex with its four Ig-like domains (D1-D4) forming contacts with Fcμ1, Fcμ2, and the JC. Ordered glycans were observed at each of the potential N-linked glycosylation sites (PNGS) on SC (four) and JC (one); while only two bases were assigned for JC Asn53, the map density suggested that this glycan extended further and may contact residues in JC_W1_ (Supplementary Fig. 3a-f). Although HC Tp residue Asn435 is a conserved PNGS observed in at least one human SIgM structure^26^, evidence for glycans at this site in *ap* SFcμJ was not apparent in the map.

### The FcμJ core of avian SIgM

The FcμJ core, comprising the five Fcμ subunits and one JC, adopts hexagonal symmetry with the JC occupying the gap between Fcμ1 and Fcμ5. The central angles between contacting subunits measured approximately 60° (Fig. 2a), comparable to the FcμJ core in human SIgM structures^26,27^. Together, the Fcμ subunits are relatively planar, with each intersecting a common central plane. However, we observed each Fcμ subunit deviating uniquely from the plane, occupying a distinct position relative to the other subunits (Fig. 2b). In reported avian IgA structures, Fcα subunit deviations from the central plane were quantified using two measurements, termed *twist* and *tilt*, and were predicted to influence the positions of Fabs^43^. Accordingly, we measured twist and tilt values of Fc subunits in *ap* SFcμJ and compared them to the values measured in human IgM and avian IgA structures. All Fcμ subunit twist values were negative in *ap* SFcμJ, ranging from –8.4° (Fcμ3) to –1.5° (Fcμ5), with more atoms in each subunit’s chain A positioned above the central plane compared to chain B (Fig. 2b, Supplementary Fig. 4). In *ap* SFcμJ, Fcμ subunits’ tilt values (i.e. projection out of the central plane) transitioned sequentially from projecting toward the front face (+4.9° for Fcμ1) to projecting toward the back face (−13.2° for Fcμ5) (Fig. 2b, Supplementary Fig. 4). In human SFcμJ, all Fcμ subunits exhibited positive twist values (i.e. for each subunit, fewer atoms in chain A were positioned above the central plane compared to chain B) which differed from that observed in *ap* SFcμJ; however, tilt values followed a similar pattern in human SFcμJ as that observed in the avian complex (Supplementary Fig. 4). Comparison of *ap* SFcμJ to avian IgA structures revealed Fc subunit projection out of the front and back faces to be closely matched, with tilt of Fcμ5 in *ap* SFcμJ being more similar to the equivalent subunit in *ap* SFcαJ (Fcα2) than to Fcμ5 in human SFcμJ (Supplementary Fig. 4), suggesting species-specific constraints on subunit positioning across isotypes. Nevertheless, the similar directionality of twist values (i.e. all positive or all negative) and tilt values (i.e. sequentially transitioning from projection toward the front face to projection toward the front face) across all five Fcμ subunits within each IgM complex suggests that the spatial relationship among subunits within the IgM pentameric core is conserved across species, possibly reflecting functional constraints on the pentameric state.

**Figure 2.**
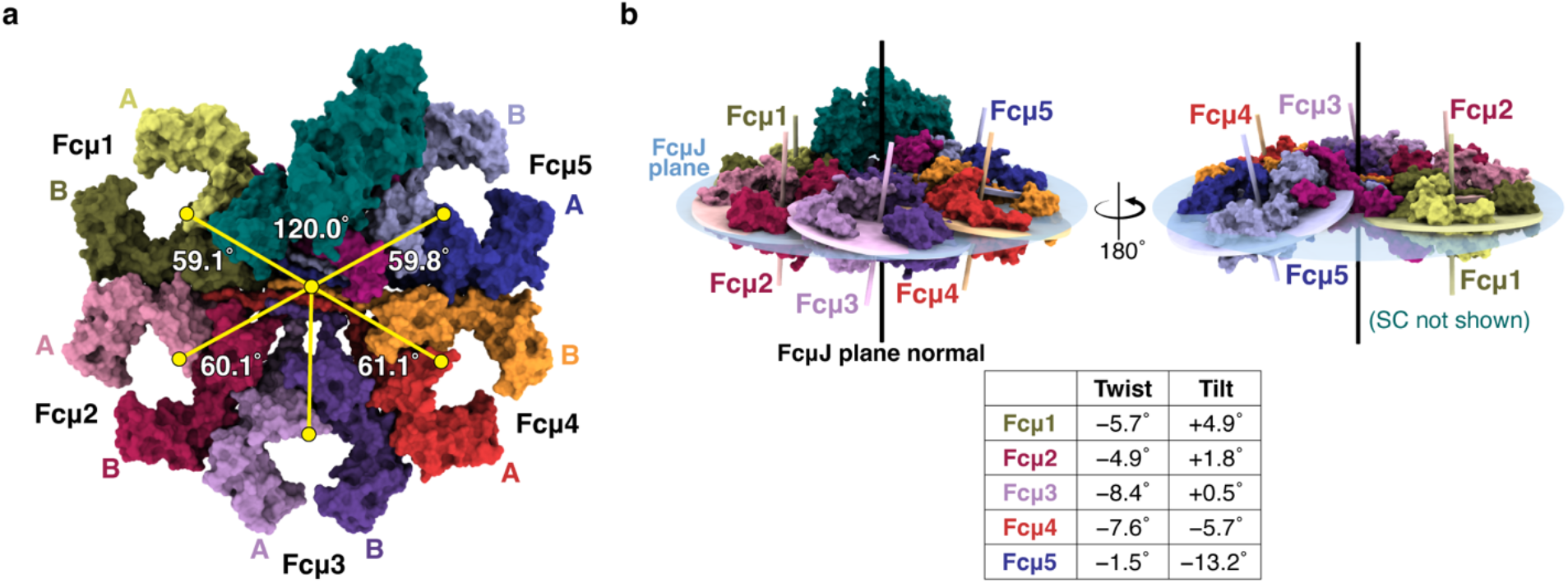
*ap* SFcμJ core geometry. (**a**) Surface representation of *ap* SFcμJ front face; HC and JC are colored as in (Fig. 1) and SC is colored dark teal. FcμJ and Fcμ subunit centroids are shown as yellow circles and central angles formed between adjacent Fcμ subunits are reported in white text. (**b**) The *ap* SFcμJ structure shown from two views, with (left) and without (right) SC. The FcμJ plane (light blue, semi-transparent ellipse), FcμJ plane normal (black line), Fcμ subunit planes (colored ellipses), and Fcμ subunit plane normals (colored lines) are also shown. Twist and tilt values for each subunit are given in the table below.

In *ap* SFcμJ, Fcμ-Fcμ interfaces were extensive, with buried surface areas ranging from ∼780 Å^2^ to ∼824 Å^2^ (Fig. 3a, b, Supplementary Fig. 5a-f). These values were notably larger than the Fcμ-Fcμ interfaces in human SFcμJ, which ranged from ∼527 Å^2^ to ∼578 Å^2^ (Supplementary Fig. 6a). Consistent with human SIgM structures, interchain disulfide bonds, which are known to stabilize Fcμ-Fcμ interfaces, were observed at each *ap* SFcμJ Fcμ-Fcμ interface between adjacent copies of Cys286 (Fig. 3b, Supplementary Fig. 5f). While some polar interactions between interfacing residues were observed, Fcμ-Fcμ interfaces in *ap* SFcμJ appeared to be largely stabilized by hydrophobic interactions, with ∼48% of interfacing residues being hydrophobic compared to ∼32% in human SFcμJ (Fig. 3b, Supplementary Fig. 5a-e, 6a-c). Overall, the four Fcμ-Fcμ interfaces in *ap* SFcμJ were comparable, with marginal variation in contacting residues observed (Supplementary Fig. 5a-e). In contrast, the two Fcα-Fcα interfaces in avian IgA structures were reported to display differences in buried surface area and interacting residues^43^ (Supplementary Fig. 6a-c). Specifically, the *ap* SFcαJ Fcα1-Fcα3 interface buried more surface area (993 Å^2^) than any *ap* SFcμJ Fcμ-Fcμ interface, despite involving a smaller percentage of hydrophobic residues (30%), while the Fcα2-Fcα4 interface buried less surface area (587 Å^2^), but involved a greater percentage of hydrophobic residues (50%) (Supplementary Fig. 6a-c). The similarity among Fcμ-Fcμ interfaces within a given complex appears to be a conserved feature of pentameric IgM across species, distinguishing it from the asymmetric Fcα-Fcα interfaces characteristic of *ap* SFcαJ.

**Figure 3.**
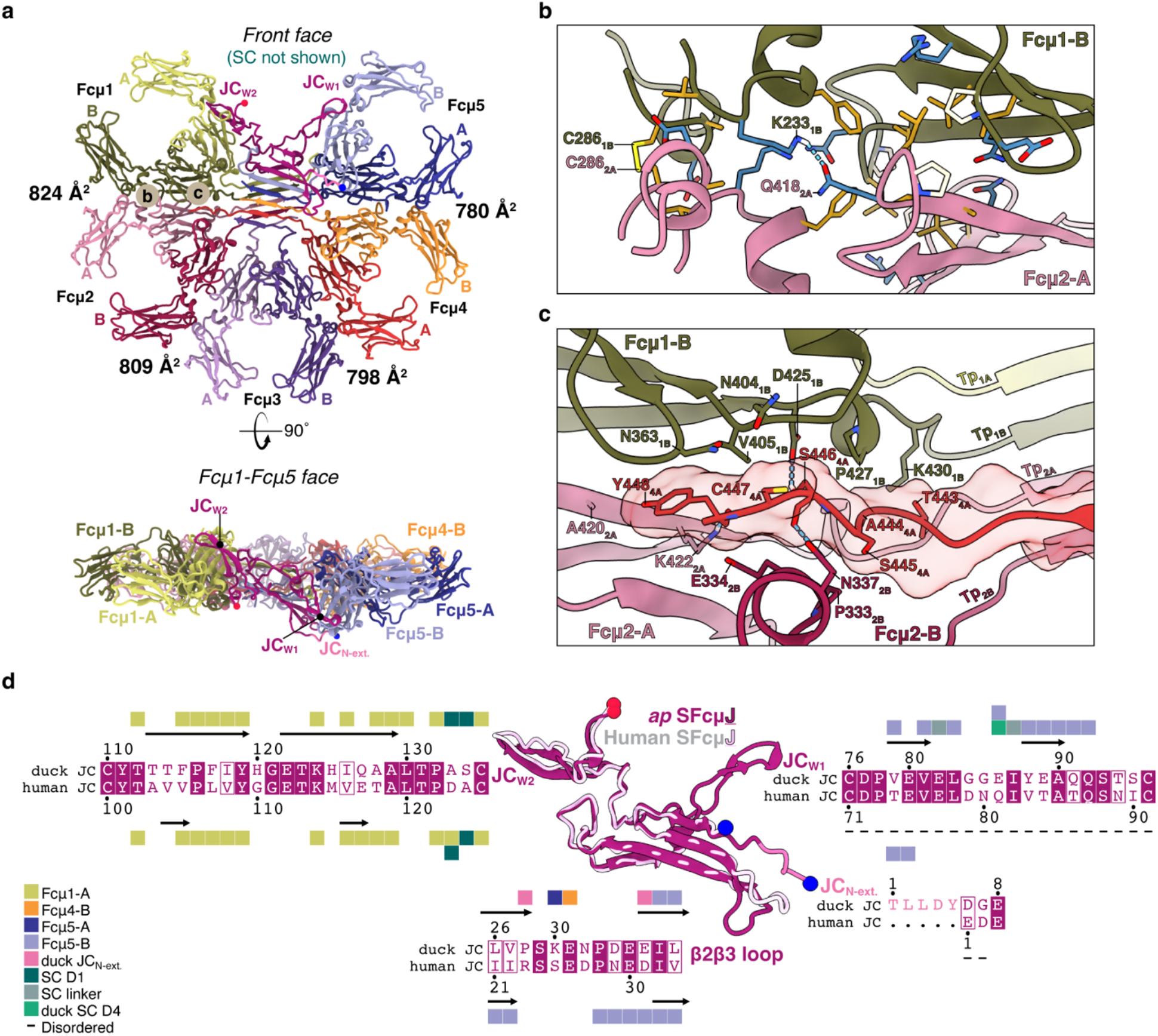
Molecular interfaces of FcμJ core in *ap* SFcμJ. (**a**) Cartoon representation of *ap* SFcμJ front and top face views; colored as in (Fig. 1) with SC removed. JC N– and C-termini are indicated by blue and red spheres, respectively; JC_W1_ and JC_W2_ are indicated. Buried surface area for Fcμ-Fcμ interfaces is shown in black text. Circled letters indicate the regions enlarged in corresponding figure panels. (**b**) Fcμ1-Fcμ2 interface shown as a cartoon with residues involved in contacts shown as sticks and colored either gold (hydrophobic), cream (neutral), or blue (hydrophilic). Residues participating in interchain disulfide or hydrogen bond are labeled. (**c**) Fcμ1-Fcμ2-Tp_4A_ interface shown as a cartoon with residues involved in contacts shown as sticks and labeled. Semi-transparent molecular surface of Tp_4A_ is also shown. (**d**) Cartoon representation of *ap* SFcμJ JC (magenta) aligned to the human SFcμJ JC (PDB 6KXS; light pink) shown in center. JC N– and C-termini are indicated by blue and red spheres, respectively; JC structural motifs are also labeled. Structure-based sequence alignments of duck and human JC motifs are shown with colored squares to indicate contacts according to key. Dashes signify disordered residues.

The *ap* SFcμJ structure also showed evidence that a subset of Tps may directly stabilize Fcμ-Fcμ interfaces in a manner distinct from other structures. The Fcμ1-Fcμ2 interface is stabilized by interactions with Tp_4A_ residues 442-448, which follow the terminal β-strand and sit in a pocket on the front face of *ap* SFcμJ between the two Fcμ subunits where they contact Fcμ1-B, Fcμ2-A, and Fcμ2-B (Fig. 3c). With each of these chains, Tp_4A_ buries surface areas of ∼194 Å^2^, ∼186 Å^2^, and ∼160 Å^2^, respectively, resulting in a cumulative Fcμ1-Fcμ2-Tp_4A_ buried surface area of ∼1365 Å^2^. Notably, we also observed additional density at this interface, which may correspond to unassigned residues from a neighboring Tp that may also contact the Tp_4A_ C-terminus (Supplementary Fig. 7a, b). Similarly, we observed unassigned density resembling one or more C-terminal Tp residues on the back face between Fcμ4 and Fcμ5 (Supplementary Fig. 7a, c). While our inspection of reported human SIgM maps^26,27^ (EMDBs 0782, 22591) show density suggesting C-terminal Tp residues contact proximally located Cμ4 residues, we did not find evidence of Tps participating in *ap* SFcμJ Fcμ-Fcμ interfaces.

In addition to the four Fcμ-Fcμ interfaces, the *ap* FcμJ core is stabilized by interfaces formed between the HCs and the JC; the JC_core_ buries the largest cumulative HC surface area (∼2119 Å^2^) compared to JC_W1_ (∼521 Å^2^) and JC_W2_ (∼698 Å^2^), forming contacts with three of the five subunits (Fcμ1, Fcμ4, and Fcμ5) (Fig. 3a, d). The JC_core_-Tp assembly is structurally similar to that observed in human SFcμJ and *ap* SFcαJ structures (RMSD of 0.943 when aligned to Cα atoms in PDB 6KXS and RMSD of 0.679 when aligned to PDB 9EC6); however we observed local structural differences between the two species, including the conformation of the loop connecting the β2 and β3 strands (β2β3 loop) (RMSD of 3.671) (Supplementary Fig. 8a), and the presence of the four-residue JC_N-ext,_ which is absent from mammalian sequences. Instead of contacting residues in Fcμ5-B Cμ4, as observed in the human SFcμJ structure, the avian β2β3 loop contacts the JC_N-ext._, which bridges the interface between the β2β3 loop and Fcμ5-B (Fig. 3d, Supplementary Fig. 8b). In *ap* SFcαJ, the β2β3 loop adopts a similar conformation (RMSD of 0.540) and the JC_N-ext._ also bridges the interface between the β2β3 loop and Fcα2-C, albeit with more contacts than observed in the *ap* SFcμJ structure (Supplementary Fig. 8b). Additionally, the JC β2β3 loop contacts one or more residues in the Cμ4/Cα4-Tp linker in Fcμ4-B/Fcα4-G in *ap* SFcμJ and avian IgA structures (Supplementary Fig. 8b).

In *ap* SFcμJ, JC_W1_ and JC_W2_ appear to play similar structural roles as observed in mammalian JC-containing structures and the avian IgA structures; however as noted, JC_W1_ (residues 76 to 96) is ordered, forming an interface with Fcμ5-B (Fig. 3d) and in contrast with human SIgM structures. JC_W1_ in *ap* SFcμJ is positioned similarly to ordered JC_W1_ in both human IgM–CD5L structures and *ap* SFcαJ, with Cα alignment yielding RMSDs of 0.803 and 0.847, respectively (Supplementary Fig. 8a). The position of JC_W2_ (residues 110 to 135) in *ap* SFcμJ is also comparable to the position in human FcμJ (RMSD of 0.857) and *ap* SFcαJ (RMSD of 0.601) (Supplementary Fig. 8a), forming contacts with Fcμ1-A and Fcα1-A, respectively.

### Avian SC shares an extensive interface with IgM

Comparison of the SC-FcμJ interface in the *ap* SFcμJ structure with human SIgM structures revealed both conserved and unique features. SC binds to the front face of both structures, with the first domain (D1) contacting residues in Fcμ1 and the JC, while the terminal domain (D4 in birds, D5 in mammals) interfaces Fcμ5 (Fig. 4a, Supplementary Fig. 9a, b). In *ap* SFcμJ, SC D1 (residues 12-121) contacts residues in Fcμ1, a linker connecting JC_W1_ and JC_W2_, the JC C-terminus, and Tp_5B_ using residues in its CDR-like loops (hereafter, CDRs) (Fig. 4b, Supplementary Fig. 9a, b). While the D1-FcμJ interfaces were mostly conserved between *ap* SFcμJ and human SIgM complexes, differences were observed at D1 CDR3 and the proximally located CCʹ loop (residues 52-55) (Supplementary Fig. 9a, b). In human SIgM structures, the CCʹ loop contacts Tp_4B_ while CDR3 contacts both Tp_5A_ and Tp_5B_ (Supplementary Fig. 10a). In *ap* SFcμJ, CDR3 also contacts Tp_5B_, however, contacts between the CCʹ loop and Tp_4B_ were not apparent (Supplementary Fig. 10b). The *ap* SFcμJ density map suggested additional contacts between CDR3 and CCʹ loop residues and other avian Tp residues, but these could not be reliably assigned (Supplementary Fig. 10b). Furthermore, when aligning human SFcμJ to the *ap* SFcμJ structure and map, avian Tp density did not correspond well to human Tp positioning, indicating that Tp_4B_ and Tp_5A_ likely occupy different positions and/or have more conformational variability in *ap* SFcμJ. When comparing Tp densities between *ap* SFcαJ and mammalian SIgA structures^43^, this difference was also observed (Supplementary Fig. 10c). Together, these observations suggest that avian Tps interface SC D1 differently than their mammalian counterparts, which may reflect species-specific differences in SC binding to both IgM and IgA.

**Figure 4.**
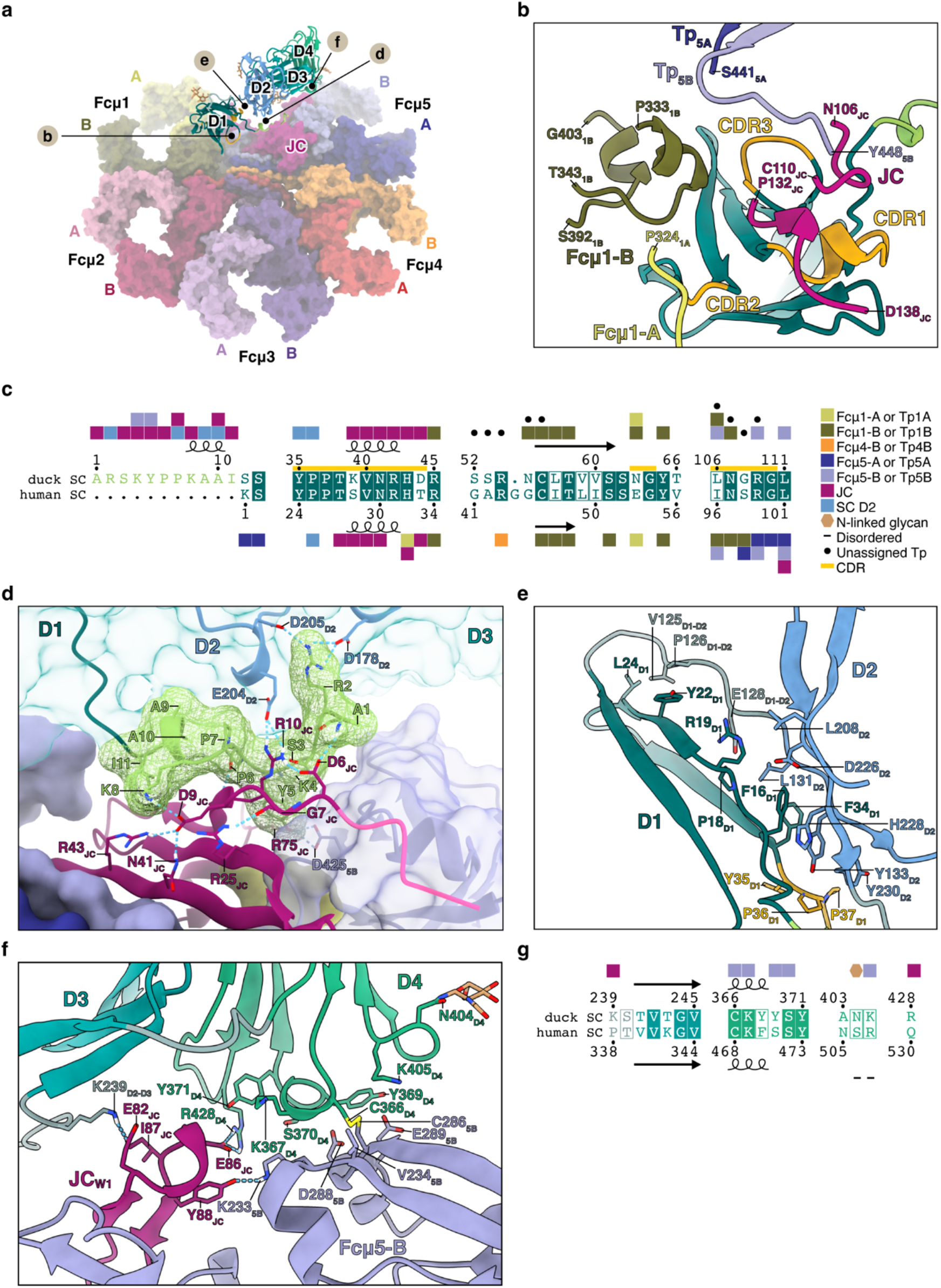
SC contacts with FcμJ core in *ap* SFcμJ. (**a**) Overall structure of *ap* SFcμJ with HCs and JC shown as surfaces and SC as a cartoon. Components are colored as in (Fig. 1). Glycans are shown as sticks. Circled letters indicate the corresponding figure panels that show enlarged views of the area. (**b**) Duck SC cartoon showing D1 interfaces with HCs and JC. CDRs are colored gold and numbered. (**c**) Structure-based sequence alignment of duck SC and human SC showing secondary structure elements. Structural motifs and residue contacts with other chains are indicated according to the key. (**d**) SC_N-ext._ residues and their contacts in *ap* SFcμJ are shown as cartoons and sticks. Molecular surfaces of SC (transparent teal), Fcμ5-B (lavender), and the SC_N-ext._ (green mesh) are shown; interacting residues and SC domains are labeled. (**e**) SC D1-D2 interface shown as a cartoon with contacting residues labeled and shown as sticks. (**f**) Duck SC D4 interface with Fcμ5-B and the JC_W1_. Contacting residues and N404-linked glycosylation are labeled and shown as sticks. (**g**) Structure-based sequence alignment of indicated duck SC D4 and human SC D5 residues. Residue contacts with other chains are indicated by key in (c). Dashes indicate disordered residues.

In *ap* SFcμJ, we also observed the SC_N-ext._, 11 N-terminal residues not conserved in mammals, bridging interactions between SC D1 and the FcμJ core. The SC_N-ext_ contributed approximately one third of the total SC D1 interface with the FcμJ core (∼499 Å^2^ out of ∼1465 Å^2^), while also forming a ∼298 Å^2^ interface with D2 (Fig. 4c, d, Supplementary Fig. 9a). The SC_N-ext._ fills a pocket that is unoccupied in human SIgM structures, where human SC D1 buries only ∼1001 Å^2^ on the FcμJ core (Supplementary Fig. 9a, b, 11a). The SC_N-ext._ was ordered in the *ap* SFcαJ structure^43^ (Supplementary Fig. 11b); comparison between *ap* SFcμJ and *ap* SFcαJ structures revealed SC_N-ext._ Arg2 forming several contacts with D2 residues in *ap* SFcμJ rather than being oriented away from D2 as observed in *ap* SFcαJ (Fig. 4c, d, Supplementary Fig. 9a, b, 11b). When comparing buried surface area on D2, the SC_N-ext._ buries an additional ∼93 Å^2^ in *ap* SFcμJ compared to *ap* SFcαJ. The remaining SC_N-ext._ residues occupied similar positions in both structures with Tyr5 bridging contacts between Fcμ5-B and the JC_core_ and subsequent residues interfacing the JC_core_ and D2 (Fig. 4c, d, Supplementary Fig. 9a, b, 11). In *ap* SFcμJ, the SC_N-ext._ forms an interface with JC of ∼400 Å^2^, which includes charged JC N-terminal residues Asp6, Glu9, and Arg10, as well as residues in the β1 and β2 strands, and is comparable to the SC_N-ext._-JC interface in *ap* SFcαJ (Fig. 4c, d, Supplementary Fig. 8b, 9b). The isotype-varied positioning of the SC_N-ext._ may stem from differences in the sizes and chemical properties of nearby Cμ4 residues; nonetheless, the structural role of the SC_N-ext._ appears to be conserved between the avian SFcμJ and SFcαJ structures.

In birds, SC D1 is connected to the remaining three domains via a linker rather than an additional domain homologous to mammalian D2. In *ap* SFcμJ, SC D1 and D2 and their linking residues form an interface (Fig. 4e) that is nearly identical to that reported in the *ap* SFcαJ structure, where this linker was shown to increase contacts between D1 and D2, resulting in higher buried hydrophobic surface area compared to mammalian SC^43^. SC D3 occupies a position comparable to D3 in *ap* SFcαJ, interfacing D2 and D4, but forming few contacts with the FcμJ core (Fig. 4f, g, Supplementary Fig. 9b).

SC D4, the terminal domain, interfaced with Fcμ5-B Cμ3 and JC_W1_ in *ap* SFcμJ (Fig. 4f, g). The interface revealed evidence for a disulfide bridge between D4 Cys366 and Fcμ5-B Cys286, hydrophobic and polar contacts between D4 BC and DE loops and Fcμ5-B, and polar and electrostatic interactions between D4 FG loop and JC_W1_ (Fig. 4f, g). This interface was similar to that observed between human SC D5 and IgM (Supplementary Fig. 9a, b); however, direct comparisons were limited by disorder in JC_W1_ and a subset of the SC D5 loops in human SIgM structures^26,27^. Comparison with the *ap* SFcαJ structure revealed a similar interface, though D4 buries a modestly larger surface area on the homologous Fcα2-C chain (∼281 Å^2^) compared to Fcμ5-B (∼223 Å^2^) in *ap* SFcμJ (Supplementary Fig. 9a, b), consistent with minor differences in the positioning of these homologous chains between complexes. Together, these data reveal that terminal SC domain interactions with pIgs are broadly conserved between birds and mammals, with the highest similarity observed between avian SC binding to avian IgM and IgA, suggesting that avian SC evolved to support comparable modes of binding to both isotypes.

### Unique avian SC structural features impact IgM binding

Similarities and differences observed at the SC-pIg interface across *ap* SFcμJ, human SIgM, and *ap* SFcαJ structures, suggested that pIgR binding involves both conserved and species-specific features. To investigate how unique avian SC structural features contribute to SC-IgM and SC-IgA interactions, we used surface plasmon resonance (SPR). SC and D1 are established soluble models for mammalian pIgR-IgM/IgA binding ^4,6,45^; as their multi-step binding process does not fit 1:1 kinetic models, we qualitatively compared concentration-matched responses and concentration-matched normalized responses for analyte-ligand pairs. To characterize avian SC binding to IgM and assess contributions from the JC_N-ext._ and SC_N-ext._, we compared responses of *ap* SC wt and an SC variant lacking the SC_N-ext._ (*ap* SC D1_ΔN-ext._-D2-D3-D4) to immobilized ligands, *ap* FcμJ and *ap* FcμJ lacking the JC_N-ext._ (*ap* FcμJ_1-4del_). The *ap* SC wt displayed concentration-dependent binding to *ap* FcμJ with a relatively moderate association and slow dissociation (Fig. 5a; Supplementary Fig. 12a). When binding to the *ap* FcμJ_1-4del_ ligand, *ap* SC wt also displayed relatively moderate association and slow dissociation (Fig. 5b; Supplementary Fig. 12b), suggesting that the JC_N-ext._ does not significantly impact SC-IgM binding or SIgM complex stability. In contrast, *ap* SC D1_ΔN-ext._-D2-D3-D4 sensorgrams displayed a marked reduction in maximum response units (RU; 4 RU at 64 nM compared to 123 RU for *ap* SC wt) as well as more rapid association and dissociation phases compared with concentration-matched *ap* SC wt responses for both *ap* FcμJ and *ap* FcμJ_1-4del_ ligands (Fig. 5a,b; Supplementary Fig. 12a, b). These results indicate weaker binding and suggest that the SC_N-ext._ contributes to avian SC-IgM binding kinetics.

**Figure 5.**
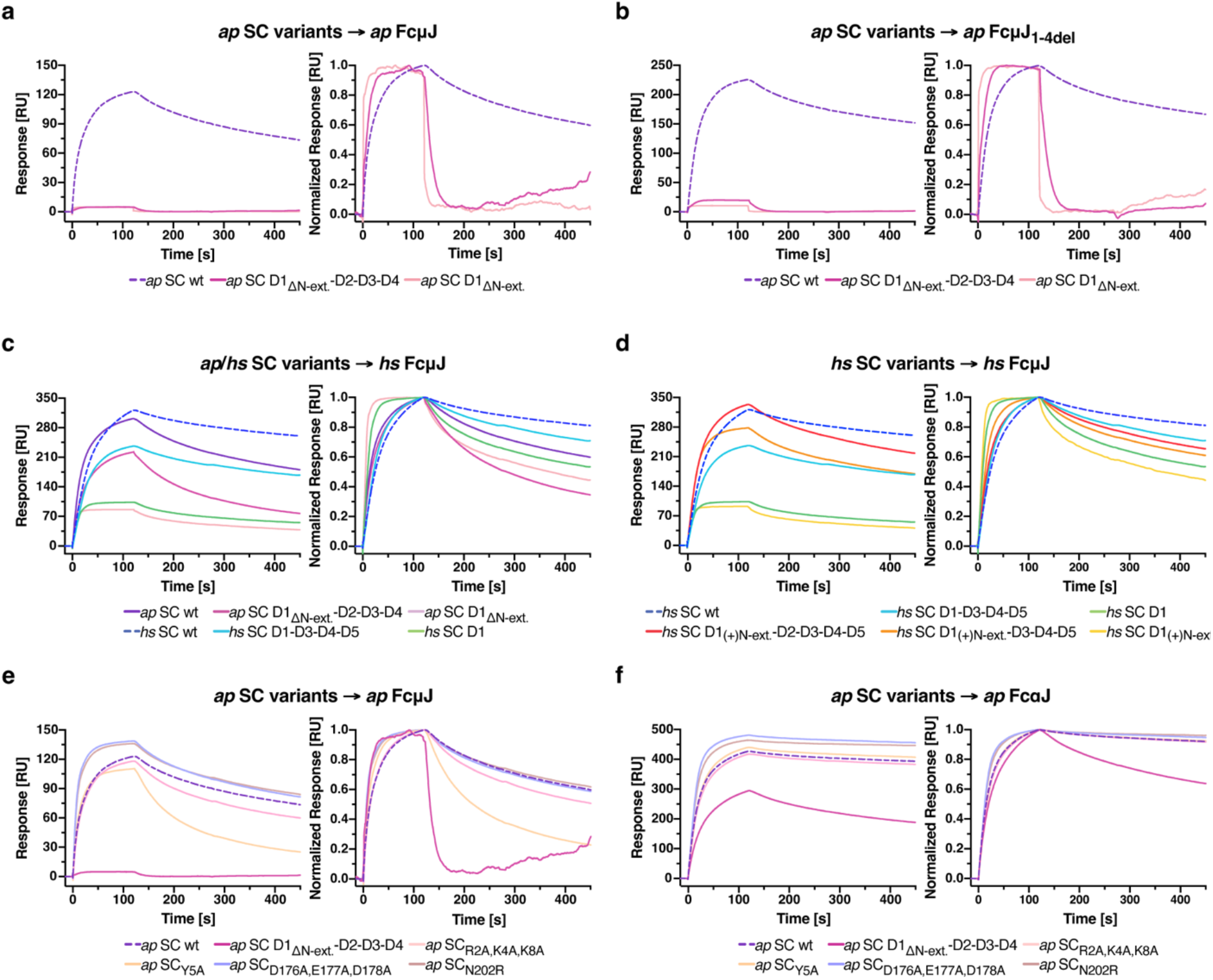
Duck and human SC variant binding to cognate and non-cognate ligands. (**a**-**f**) SPR sensorgrams showing the response (left) and normalized response (right) of 64 nM SC variant analytes binding to FcμJ or FcαJ ligands for the following: (a, e) *ap* SC variants to *ap* FcμJ, (b) *ap* SC variants to *ap* FcμJ_1-4del_, (c) *ap* SC and human (*hs*) SC variants to *hs* FcμJ, (d) *hs* SC and chimeric *hs* SC variants to *hs* FcμJ, and (f) *ap* SC variants to *ap* FcαJ. Each analyte response is colored according to the key at the bottom of the panel and dashed lines indicate cognate interactions between unmutated variants. Results are consistent with three or more replicate experiments. Sensorgrams for the full analyte concentration series of each experiment are provided in Supplementary Fig. 12-14.

To assess species-specific differences in avian SC-IgM interactions, we assayed *ap* SC variant binding to human (*hs*) FcμJ and compared to *hs* SC binding to *hs* FcμJ. The *ap* SC wt displayed concentration-dependent binding to *hs* FcμJ; however, dissociation was more rapid than observed for *hs* SC wt binding to *hs* FcμJ (Fig. 5c; Supplementary Fig. 13a, b), suggesting that despite conservation of some SC-IgM interactions across species, SC and IgM have coevolved within each species such that non-cognate pairs form less stable complexes. We also compared binding of *ap* SC D1_ΔN-ext._-D2-D3-D4 and *hs* SC with a D2 deletion (*hs* SC D1-D3-D4-D5), which share equivalent domain organization, to *hs* FcμJ.

The *ap* SC D1_ΔN-ext._-D2-D3-D4 displayed more rapid dissociation from *hs* FcμJ compared to *hs* SC D1-D3-D4-D5 (Fig. 5c; Supplementary Fig. 13a, b), but this effect was less pronounced than observed for *ap* SC D1_ΔN-ext._-D2-D3-D4 binding to *ap* FcμJ (Fig. 5a; Supplementary Fig. 12a), indicating that the structural features that support SC-IgM complex stability may vary between birds and humans, and that the avian SC_N-ext._ serves a more critical role in avian SC binding to avian IgM than to non-cognate, human IgM. Furthermore, *ap* SC D1 binding to *hs* FcμJ was comparable to *hs* SC D1 binding to *hs* FcμJ (Fig. 5c; Supplementary Fig. 13a, b), suggesting that avian and human D1 contribute similarly to *hs* FcμJ binding in the absence of additional domains. Additionally, the SC_N-ext._ may partially compensate for the absence of human-specific structural features that support SC-IgM complex stability, as *ap* SC wt dissociation from *hs* FcμJ more closely resembled that of *hs* SC D1-D3-D4-D5 than *ap* SC D1_ΔN-ext._-D2-D3-D4 (Fig. 5c; Supplementary Fig. 13a, b). Together, these results suggest that SC and IgM have coevolved species-specific structural features that collectively support complex stability, with the avian SC_N-ext._ representing a feature that has diverged between birds and mammals.

Having observed that the avian SC_N-ext._ contributes to species-specific differences in SC-IgM interactions, we sought to determine whether this motif could also influence human SC binding to *hs* FcμJ. We compared *hs* FcμJ binding of three human SC variants (i.e. *hs* SC wt, *hs* SC D1-D3-D4-D5, and *hs* SC D1) to chimeric human SC variants containing the duck SC_N-ext._ (i.e. *hs* SC D1_(+)N-ext._-D2-D3-D4-D5, *hs* SC D1_(+)N-ext._-D3-D4-D5, and *hs* SC D1_(+)N-ext._). In all cases, the addition of the SC_N-ext_ to human SC variants resulted in more rapid dissociation from *hs* FcμJ compared to equivalent variants lacking the motif (Fig. 5d; Supplementary Fig. 13b, c). These results suggest that the avian SC_N-ext._ can influence human SC-IgM interactions, but that the effects on complex formation and stability differ from those in avian SC-IgM interactions which is consistent with the species-specific coevolution of pIgR and IgM.

The SC_N-ext._ contributions to cognate SC-IgM binding appear more pronounced than contributions to SC-IgA binding, which we recently reported^43^ and repeated for this study (Fig. 5e, f; Supplementary Fig. 14a, b). Specifically, when comparing *ap* SC wt binding to *ap* FcμJ and *ap* FcαJ ligands, a more rapid dissociation phase was observed for the *ap* FcμJ ligand. However, removal of the SC_N-ext._ had a more pronounced effect on *ap* FcμJ binding, with *ap* SC D1_ΔN-ext._-D2-D3-D4 exhibiting a 95% reduction in maximum RU for *ap* FcμJ compared to a 30% reduction for *ap* FcαJ ligand (Fig. 5e, f; Supplementary Fig. 14a, b). Together, these results point toward isotype-dependent differences in SC_N-ext._ contributions to SC ligand binding, with the SC_N-ext._ likely to play a more prominent role in the avian SC-IgM interaction. Furthermore, the similar positioning of the SC_N-ext._ in *ap* SFcμJ and *ap* SFcαJ structures (Fig. 4d, Supplementary Fig. 9b, 11b), despite marked isotype-dependent differences in binding kinetics, suggests that the SC_N-ext._ may contribute differently to the initial pIgR-IgM and pIgR-IgA interactions compared to those observed in *ap* SFcαJ and *ap* SFcμJ structures when binding is complete and SC is covalently bound.

Considering observed differences in SC_N-ext._ buried surface area on *ap* FcμJ and *ap* FcαJ complexes, including contacts with SC Arg2 (Fig. 4c, d, Supplementary Fig. 9a, b, 11b), we sought to further investigate the role of SC_N-ext._ residues and their proximal interactions on SC-IgM and SC-IgA binding. To accomplish this, we generated two SC variants with mutations in the SC_N-ext._ (*ap* SC_R2A,K4A,K8A_ and *ap* SC_Y5A_) and tested their binding to *ap* FcμJ and *ap* FcαJ. Compared with concentration-matched *ap* SC wt responses, *ap* SC_R2A,K4A,K8A_ sensorgrams exhibited no marked change in the association phase or maximum RU, yet did exhibit marginally more rapid dissociation (Fig. 5e, f; Supplementary Fig. 14a, b), suggesting that one or more charged residues in the SC_N-ext._ may promote SC-IgM complex stability. In contrast, binding between *ap* SC_R2A,K4A,K8A_ and *ap* FcαJ was comparable to *ap* SC wt (Fig. 5e, f; Supplementary Fig. 14a, b). Isotype-dependent differences in binding profiles were also observed with *ap* SC_Y5A_, which displayed more rapid association and dissociation phases for *ap* FcμJ ligand binding compared to *ap* SC wt (Fig. 5e, f; Supplementary Fig. 14a, b). Moreover, sensorgrams for *ap* SC_Y5A_ binding to *ap* FcαJ were comparable to *ap* SC wt binding (Fig. 5e, f; Supplementary Fig. 14a, b), suggesting that Tyr5 uniquely supports pIgR-IgM binding compared to pIgR-IgA binding. Mutation of D2 residues Asp176, Glu177, Asp178, and Asn202 (*ap* SC_D176A,E177A,D178A_ and *ap* SC_N202R_, respectively), which contact the SC_N-ext._, had only minor effects on binding that were comparable for both IgM and IgA isotypes (Fig. 5e, f; Supplementary Fig. 14a, b), suggesting that intrachain interactions between the SC_N-ext._ and D2 loops do not contribute significantly to the isotype-dependent differences in SC binding. Together, these results indicate that SC_N-ext._ residues contribute differently to pIgR-IgM and pIgR-IgA binding, with one or more charged residues Arg2, Lys4, and Lys8 supporting SC-IgM complex stability and Tyr5 supporting both pIgR-IgM association and SC-IgM stability.

## DISCUSSION

The cryo-EM structure of *ap* SFcμJ reported here provides high-resolution information on JC-dependent polymeric IgM from a non-mammalian species. The overall quaternary organization of *ap* SFcμJ is globally similar to reported human SIgM structures^26,27^, with a pentameric arrangement of Fcμ subunits, a central JC-Tp assembly, and SC binding asymmetrically to the front face of the complex. The conservation of this pentameric organization over 300 million years of evolutionary divergence between birds and mammals suggests that this polymeric state and its associated effector functions are under strong selective pressure. However, comparison of avian and human SIgM structures also revealed notable species-specific differences which may reflect the distinct immunological landscape of birds.

A noteworthy species-specific feature of the *ap* SFcμJ structure is structured JC_W1_, which is disordered in previously reported human SIgM structures^26,27^. In humans, JC_W1_ is ordered when IgM is bound by CD5L^28,29^, adopting a conformation that is comparable to that observed in mammalian SIgA structures^37–39^. In ducks, JC_W1_ is ordered in the context of IgA, SIgA, and SIgM, suggesting that JC_W1_ ordering is an intrinsic feature of the avian JC, in contrast to being promoted by a binding partner in mammals. Whether an avian homolog of CD5L exists and engages duck IgM in a comparable manner has not been established, though sequence alignments reveal conservation of some HC and JC residues that contact CD5L in humans (Supplementary Fig. 8b, 15). Additionally, in ducks, and likely other bird species, it is possible that SC may stabilize JC_W1_ in IgM, and, in its absence, JC_W1_ may be disordered. The functional implications of constitutive JC_W1_ stability in birds remain to be determined, though it may reflect divergence in JC-dependent binding partner interactions across species.

The distinct FcμJ core geometry, characterized by observed differences in tilt and twist in duck versus human SIgM structures is likely to have functional consequences. Previous modeling of mammalian IgA demonstrated that the relative positioning of Fcα subunits influences the range of conformations accessible to Fabs^38^. Additionally, human IgM Fabs have been shown to adopt preferred orientations relative to the IgM plane^44^. The opposite twist direction observed in duck SIgM compared to human SIgM suggests that IgM Fab positioning may differ between the two species, with potential consequences for antigen engagement as well as the accessibility of effector sites in the FcμJ core. Beyond Fab positioning, the opposing, yet consistent, directionality of twist across all five Fcμ subunits between the two species may reflect species-specific differences in polymeric IgM assembly, such as the order in which Fcμ monomers or JC are incorporated into the growing polymer. While polymeric Ig assembly mechanisms remain poorly understood, the directional difference in twist observed here provides a structural basis for future investigations into pentameric IgM assembly across species.

At a broader comparative level, the differences between avian and mammalian IgA are far more pronounced than those between avian and mammalian IgM, encompassing the dominant polymeric state (tetrameric duck IgA vs. dimeric mammalian IgA) and heavy chain domain organization (four constant domains in ducks vs. three constant domains in mammals). This pattern is consistent with the hypothesis that the pentameric quaternary organization of IgM has been subject to significant constraints across vertebrate evolution as it must serve both circulatory and mucosal roles. Given the strength of this constraint, we expect this *ap* SFcμJ structure to be broadly representative of SIgM across avian species. By contrast, SIgA, which functions primarily at mucosal surfaces may have been subject to different constraints, permitting diversity in polymeric state and domain organization in order to meet species-specific immune needs as a specialized mucosal antibody.

The *ap* SFcμJ structure, having SC bound, supports the existence of IgM mucosal transport via pIgR and functional relevance in avian mucosa. Despite differences in SC domain organization, our data reveal that the IgM-bound conformation of SC is structurally conserved between ducks and humans, as duck SC D1-D2-D3-D4 is superimposable with human SC D1-D3-D4-D5. However, the duck SC-IgM interface is expanded relative to the human SC-IgM interface, which is attributable to both the SC_N-ext._ and ordered JC_W1_ contacts in the *ap* SFcμJ. Moreover, SPR data indicate a dominant role for SC_N-ext._ in SC-IgM interactions compared to IgA, suggesting selective pressure to maintain mucosal IgM delivery in birds.

While our structure represents the secretory form of IgM populating mucosal surfaces in ducks, we anticipate that it also serves as a reliable model for the structure of duck circulatory IgM. Comparisons of human IgM and SIgM structures have shown that the FcμJ core is superimposable between the two forms, indicating that SC does not induce marked conformational changes^44^. Therefore, the *ap* SFcμJ structure can provide insight to mucosal, as well as circulatory, functions of IgM.

One major effector function of circulatory IgM in mammals is complement fixation through the classical pathway, where antigen-bound IgM undergoes a conformational change which exposes C1q binding sites on the Cμ3 domains^3,46,47^. Consistent with a conserved role for IgM in complement-mediated defense, duck IgM has been shown to fix complement^12,48^. IgY (an avian ortholog to mammalian IgG) has also been shown to fix complement^12,48,49^. Ducks also produce a second IgY isoform, IgY(ΔFc), which lacks the two terminal constant domains and consequently is deficient in terms of secondary effector functions, such as complement fixation. Typically, the truncated form is present in larger quantities than the full-length form, with the amount of IgY(ΔFc) continuing to increase over the course of a prolonged immune response^12^; therefore, in this context, IgM may serve as a critical contributor to complement-mediated clearance in ducks. While a high-resolution structure of the C1q-IgM has not yet been published, subtomogram averaging of human IgM-C1q complexes has confirmed that C1q interfaces the BC and FG loops in pentameric Cμ3, with the latter being critical for binding^46^. Sequence alignment of mammalian and duck IgM reveals partial conservation of residues in these loops (Supplementary Fig. 15). Combined with the conserved overall global organization between duck and human IgM and reported capacity of duck IgM to fix complement, duck IgM is likely to engage complement similarly.

Fc Receptor (FcR) mediated effector functions represent another major role of IgM in mammals; in addition to the pIgR, FcμR and Fcα/μR can also engage IgM. Despite moderate conservation of the human FcμR-contacting IgM residues at the corresponding positions in duck sequences (Supplementary Fig. 15), FcμR has not been identified in birds. Furthermore, while the duck genome appears to contain a gene encoding Fcα/μR (unpublished data), detailed comparisons between avian and mammalian Fcα/μR remain outstanding. Overall, FcR biology in ducks is poorly understood; an IgY FcR has yet to be identified and, despite conservation of most contacting residues, duck IgY does not bind the chicken FcγR, CHIR-AB1^12^. Together, these observations suggest that canonical Fc-mediated effector functions are likely to differ substantially in ducks compared to mammals and even other avian species. This divergence further highlights the importance of directly characterizing avian IgM effector functions and raises the possibility that complement-mediated functions play a proportionally greater role in duck IgM immune responses than observed in mammals.

The *ap* SFcμJ structure reported here expands the comparative structural framework for polymeric immunoglobulin organization across vertebrates, demonstrating that while the pentameric state and general SC binding mode are conserved features of SIgM, avian-specific features likely reflect both the coevolutionary history of duck Ig components and the distinct immunological landscape of birds. Notably, birds evolved rapidly relative to mammals, resulting in increased divergence between bird species, especially among immune genes (e.g. genes encoding Igs and their associated receptors) as duck and chicken immune-relevant genes reportedly display lower sequence conservation than non-immune genes^11^. Therefore, while the features observed here provide broader context for how the adaptive immune system evolved among vertebrates, they may not be uniformly observed across all avian species. Future polymeric IgM structures from additional avian species, combined with characterization of avian IgM effector functions, including FcR-dependent effector functions, will be essential to determine which features are broadly conserved across birds and which reflect specialization within the duck lineage.

## MATERIALS AND METHODS

### Construct design

Protein sequences for the *Anas platyrhynchos* (*ap*; mallard duck) IgM heavy chain (HC) constant region (GenBank: CAC43061.1) and *Homo sapiens* (*hs*; human) IgM HC constant region (UniProt: P01871) were obtained from the NCBI database. The sequences encoding the *Anas platyrhynchos* IgY (PIR: B465529) signal peptide (MSPRPHAFALLLLLAAVPGLRA)^50^, an N-terminal hexa-histidine tag, and mallard duck IgM HC residues 111 to 448 (Cμ2-Cμ3-Cμ4-Tp) were fused, codon optimized for human cell expression, synthesized (Integrated DNA Technologies, Inc.), and cloned into mammalian expression vector pD2610-v1 (Atum). Human IgM HC constructs were generated similarly after fusing the sequences encoding the tPA signal peptide, an N-terminal hexa-histidine tag, and human IgM HC residues 342 to 576 (Cμ3-Cμ4-Tp).

The following constructs were generated as previously described^43^: mallard duck IgA HC, mallard duck JC, mallard duck SC, *ap* SC wt, *ap* SC D1_ΔN-ext._-D2-D3-D4, *ap* SC D1_ΔN-ext._, human (*hs*) JC, *hs* SC wt, *hs* SC D1-D3-D4-D5, *hs* SC D1, *hs* SC D1_(+)N-ext._-D2-D3-D4-D5, *hs* SC D1_(+)N-ext._-D3-D4-D5, and *hs* SC D1_(+)N-ext._. Additional constructs, *ap* SC_R2A,K4A,K8A_, *ap* SC_Y5A_, *ap* SC_D176A,E177A,D178A_, and *ap* SC_N202R_, were generated similarly.

Amino acid sequences for all constructs are provided in Supplementary Table 2.

### Protein expression and purification

The *ap* FcμJ (mallard duck IgM HC and JC), *ap* SFcμJ (mallard duck IgM HC, JC, and SC) were transiently transfected into HEK Expi293F cells (Gibco: A14527) using previously reported approaches^43^. Complexes were purified from supernatants via affinity chromatography using Ni-NTA Agarose (Qiagen) and size exclusion chromatography (Superose 6 Increase 10/300, Cytiva). Other proteins and complexes, including *hs* FcμJ (human IgM HC and JC), *ap* FcαJ (mallard duck IgA HC and JC), *ap* SFcαJ (mallard duck IgA HC, JC, and SC), and *ap* SC variants were expressed and purified as previously described^43^.

### Cryo-EM grid preparation and data collection

UltrAuFoil R1.2/1.3 300 mesh gold grids (Quantifoil) were glow discharged in a PELCO easiGlow (Ted Pella Inc.) for 60 s at 25 mA current. Using a Vitrobot Mark IV (Thermo Fisher Scientific), three applications of 3 µL of *ap* SFcμJ at 0.1 mg/mL were applied to each grid at 4°C and 100% relative humidity. Each application was followed by a 10 s wait and 2 s blot time. Movies were collected at Purdue University on a Titan Krios G4 (Thermo Fisher Scientific) operating at 300 kV and equipped with a post-GIF K3 direct electron detector (Gatan). 3,631 movies were collected with an untilted stage using EPU (Thermo Fisher Scientific) and 3,632 movies were collected with a stage tilted at 30° using Leginon^51,52^. Collections were performed at 81,000x magnification in super resolution mode with total exposure time of 3.21 s, total dose of 57.79 electrons/Å^2^, raw pixel size of 0.527 Å/pixel, and 40 frames per movie.

### Cryo-EM data processing

All data processing was performed in cryoSPARC^53^. Raw movie frames were motion corrected with an output F-crop factor of 1/2 and contrast transfer function (CTF) corrected. Micrographs were screened based on CTF fit, ice thickness, and presence of contamination using the Manually Curate Exposures module to remove poor-quality micrographs, resulting in 3,342 untilted and 3,599 tilted micrographs. The Blob picker module was used to pick particles with a minimum and maximum particle diameter of 150 Å and 300 Å from the untilted micrographs. Picked particles were subjected to 2D Classification to generate templates for template-based particle picking. The Template picker module was used to pick particles from untilted and stage-tilted datasets with a diameter of 230 Å and minimum particle separation of 0.2 diameters. The 12.1 M picked particles underwent several rounds of 2D Classification, resulting in 507,183 particles, which were used for initial model generation and refinement using Ab-Initio Reconstruction and Non-uniform Refinement, respectively. The final refinement generated a map using 431,415 particles with an overall resolution of 3.37 Å (Fourier shell correlation (FSC) = 0.143).

### Model building, refinement, and validation

trRosetta^54^ was used to generate a *de novo* model of mallard duck IgM HC Cμ3 and Cμ4 domains. Individual domains were docked into the cryo-EM map using UCSF ChimeraX 1.8^55^. Initial inspection of the docked model revealed some loop regions fit poorly into the density; these regions were deleted from the model and then manually rebuilt using Coot 0.9.8.1^56^. IgM tailpieces were manually built using Coot while referencing previously published polymeric antibody structures^43^. The JC and SC from the mallard duck SFcαJ structure (PDB 9EC6) were docked into the cryo-EM map using UCSF ChimeraX 1.8. The entire structure was subjected to real space refinement using Coot, Phenix^57^, and ISOLDE^58^.

### Structural analysis

Geometric relationships between Fcμ subunits in *ap* SFcμJ (residues 217-426) and human SFcμJ (PDB 6KXS; residues 345-544) were determined as previously described for duck IgA^43^. Axes, planes, and centroids for the fixed coordinate system were defined using only Fcμ1, Fcμ2, Fcμ4, and Fcμ5. Twist and tilt measurements for each Fcμ subunit were determined as previously described^43^.

Buried surface area at molecular interfaces was determined using PISA^59^. Interchain contacts and molecular interfaces were determined using ChimeraX and defined as having a center-center distance less than or equal to 3.2 Å. The following structures were used for interface comparisons: human SFcμJ (PDB 6KXS), human FcμJ-CD5L (PDBs 8R83, 8R84, 8WYR, 8WYS), human FcμJ-FcμR (PDBs 7YTC, 7YTE, 7YTG, 8BPE, 8BPF, 8BPG), duck SFcαJ (PDB 9EC6).

The “super” command in PyMOL^60^ was used to calculate RMSD between Cα atoms in *ap* SFcμJ (JC β2β3 loop residues 26-38; JC_W1_ residues 76-96; JC_W2_ residues 110-135), *ap* SFcαJ (PDB 9EC6: β2β3 loop residues 26-38; JC_W1_ residues 76-96; JC_W2_ residues 110-135), human SFcμJ (PDB 6KXS; JC β2β3 loop residues 26-38; JC_W2_ residues 110-135), and human FcμJ-CD5L (PDB 8WYR; JC_W2_ residues 110-135).

### Sequence alignments

Sequence alignments were generated using ClustalOmega^61^. The alignment figure panels were generated with ESPript^62,63^ and reproduced in Adobe Illustrator.

### Surface plasmon resonance

Surface plasmon resonance (SPR) binding studies were performed using a Sierra SPR-32 Pro (Bruker) operating at 25°C. Ligands were diluted in sodium acetate buffer of either pH 4.0 (*ap* FcμJ, *ap* FcμJ_1-4del_, and *ap* FcαJ) or pH 4.5 (*hs* FcμJ) and immobilized on a High Capacity Amine Sensor (Bruker) using an amine coupling kit (Bruker). Each ligand was immobilized in a separate flow channel with at least one mock-coupled spot per channel to serve as a reference surface.

A two-fold dilution series of each analyte was prepared using HBS-EP+ buffer (10 mM HEPES, 150 mM NaCl, 3 mM EDTA, 0.05% (v/v) Tween 20) starting at 512 nM. Analytes were injected in HBS-EP+ buffer at a flow rate of 50 µL/min, with a 120 s association phase and a 320 s dissociation phase. Each injection was followed by a regeneration step (2.5 M MgCl_2_, flow rate of 25 µL/min, 60 s contact time) and two wash steps (HBS-EP+, 60 s contact time each).

Sensorgrams were reference-surface and buffer subtracted in Bruker Sierra Analyzer 3 software and then exported and replotted in GraphPad Prism 10.5.0 for macOS. Normalization of the respective sensorgrams for concentration-matched samples were carried out in Microsoft Excel by dividing each RU value by the maximum measured response of the respective analyte and then replotted in GraphPad Prism 10.5.0 for macOS. Sensorgrams shown in figures are representative of at least three replicate experiments.

### Figures

Figures were prepared using UCSF ChimeraX and Adobe Illustrator.

## ACKNOWLEDGEMENTS

The cryo-EM data collection was performed at the Purdue Cryo-EM Facility with the assistance of Thomas Klose and Frank Vago. Asta Simonovic and Sarah Leonard assisted with protein expression and purification. The authors thank Profs. Wilfred van der Donk, Nicholas Wu, Angad Mehta, and Jenna Guthmiller and all members of the Stadtmueller Lab for insightful discussions; we also thank Sarah Leonard for assistance with SPR instrumentation.

## AUTHOR CONTRIBUTIONS

The study was conceived by BMS and RMS; experiments were conducted by RMS; QL assisted with cryo-EM data collection and processing; all authors contributed to data analysis and manuscript writing.

## DATA AND MATERIALS AVAILABILITY

The Cryo-EM density map has been deposited in the Electron Microscopy Data Bank (www.ebi.ac.uk/emdb) and the refined model has been deposited in the Protein Data Bank (www.rcsb.org).

## Supplementary Materials for

**Supplementary Figure 1.**
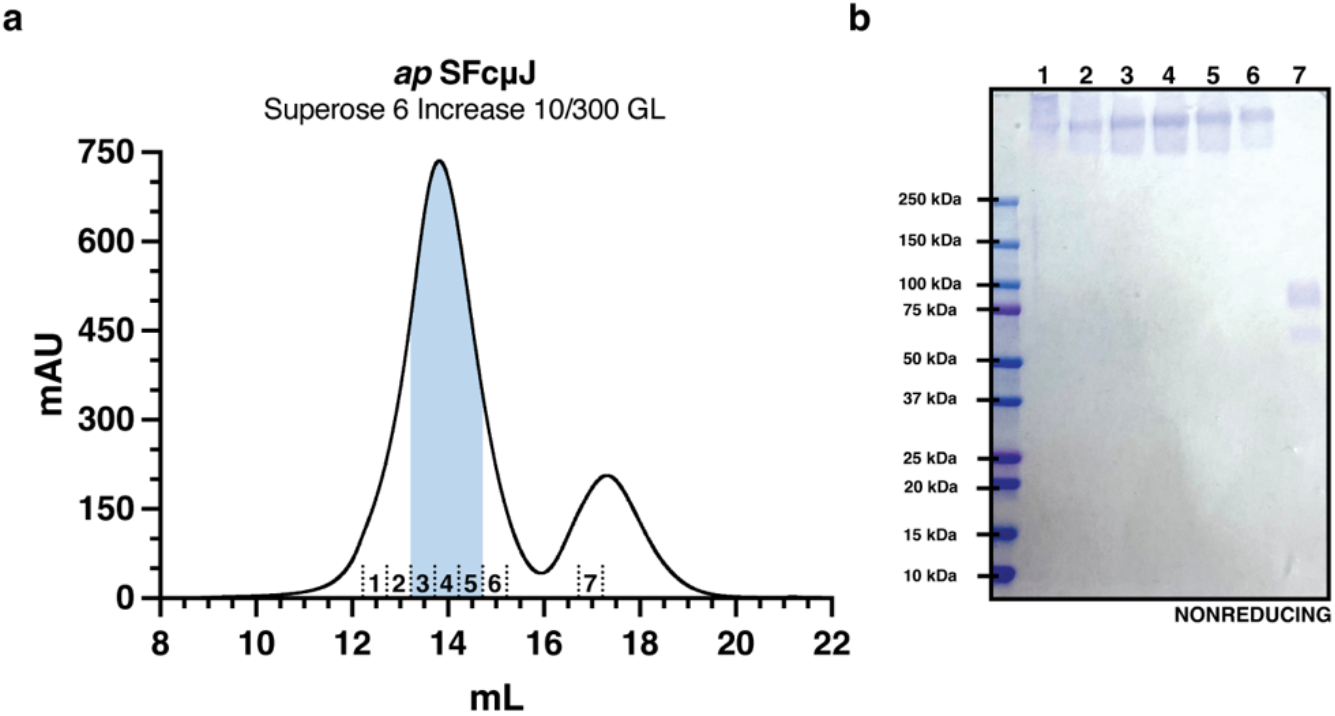
Purification of *ap* SFcμJ. (**a**) Size exclusion chromatography purification of *ap* SFcμJ after Ni-NTA affinity purification and (**b**) nonreducing SDS-PAGE gel of indicated fractions. Fractions pooled for cryo-EM imaging are indicated by blue shading in (a).

**Supplementary Figure 2.**
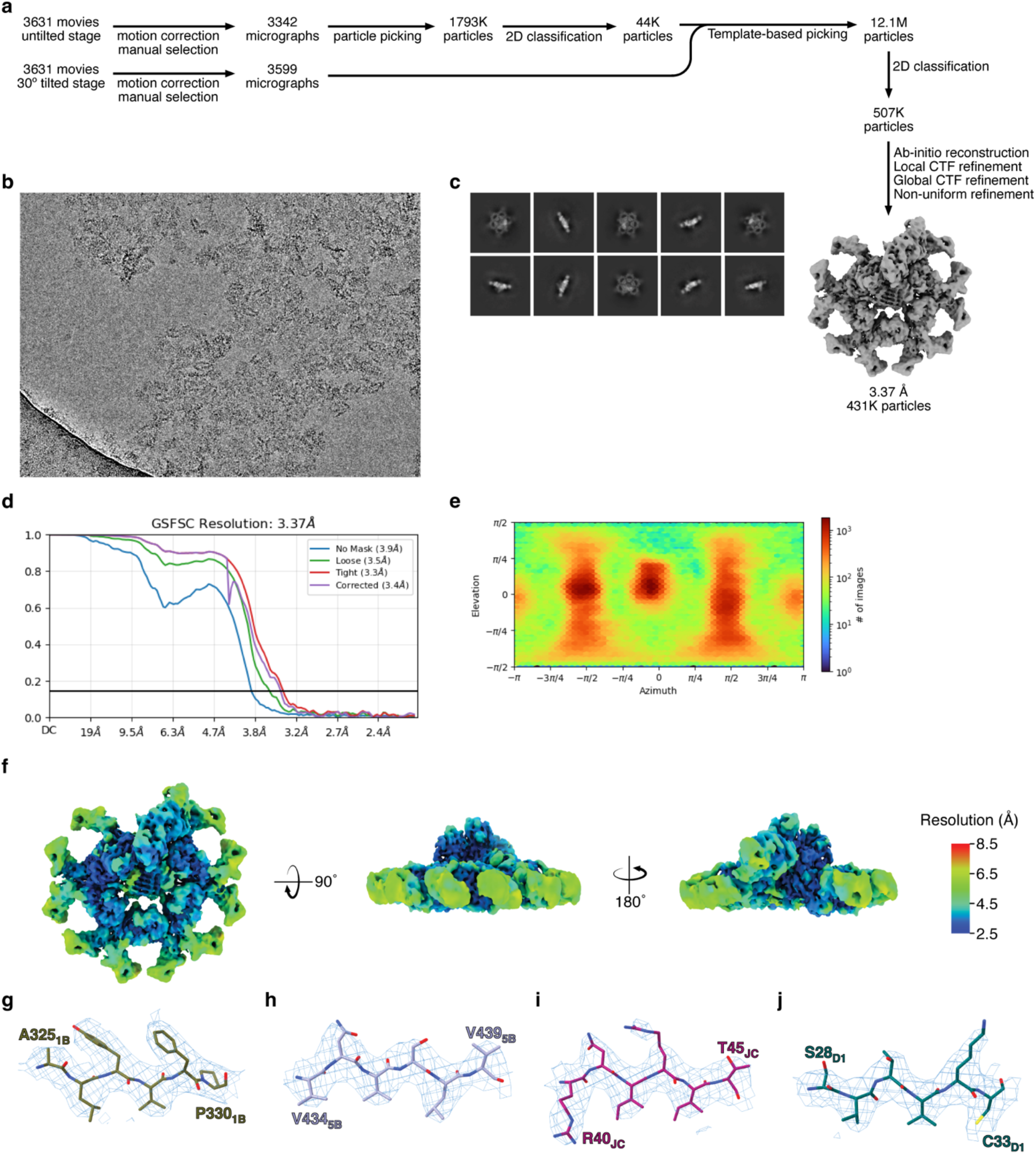
Cryo-EM data collection and cryoSPARC processing pipeline for *ap* SFcμJ. (**a**) Schematic summary of the *ap* SFcμJ cryo-EM data processing pipeline in cryoSPARC. (**b**) Representative micrograph of *ap* SFcμJ. (**c**) Representative 2D class averages. (**d**) The FSC curve for the final reconstruction with reported resolution at FSC=0.143 shown by the black horizontal line. (**e**) Viewing direction distribution plot of particles used in final refinement. (**f**) Local resolution map calculated by cryoSPARC and colored in ChimeraX. (**g**-**j**) Cryo-EM map density surrounding residues 325 to 330 in Fcμ1-B Cμ4 (g), residues 434 to 439 in Tp_5B_ (h), residues 40 to 45 in JC_core_ (i), and residues 28 to 33 in SC (j) is contoured to threshold of 0.17 and carved at 2 Å.

**Supplementary Figure 3.**
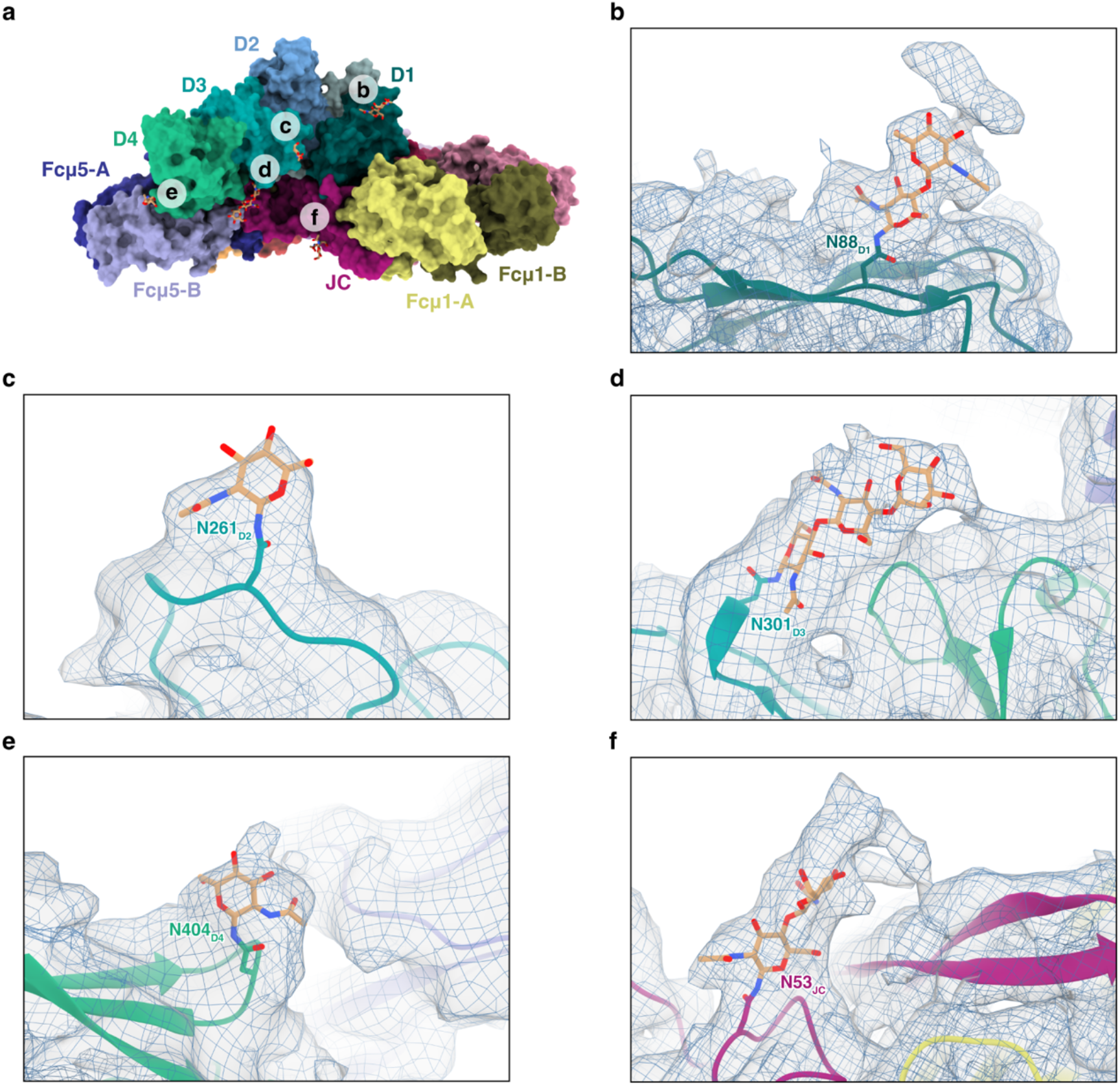
N-linked glycosylation modeled in *ap* SFcμJ cryo-EM map. (**a**) Top view of *ap* SFcμJ shown as surface representation colored as in (Fig. 1) with N-linked glycans shown as sticks. Circled letters indicate the corresponding figure panels that show enlarged views of the area. (**b**-**f**) Focused view of N-linked glycans at Asn88 (b), Asn261 (c), Asn301 (d), and Asn404 (e) in SC and Asn53 (f) in JC. Each chain is shown as a cartoon representation with the Asn side chain and glycans shown as sticks. Cryo-EM density is shown using blue mesh and light grey transparent surface at a threshold of 0.05.

**Supplementary Figure 4.**
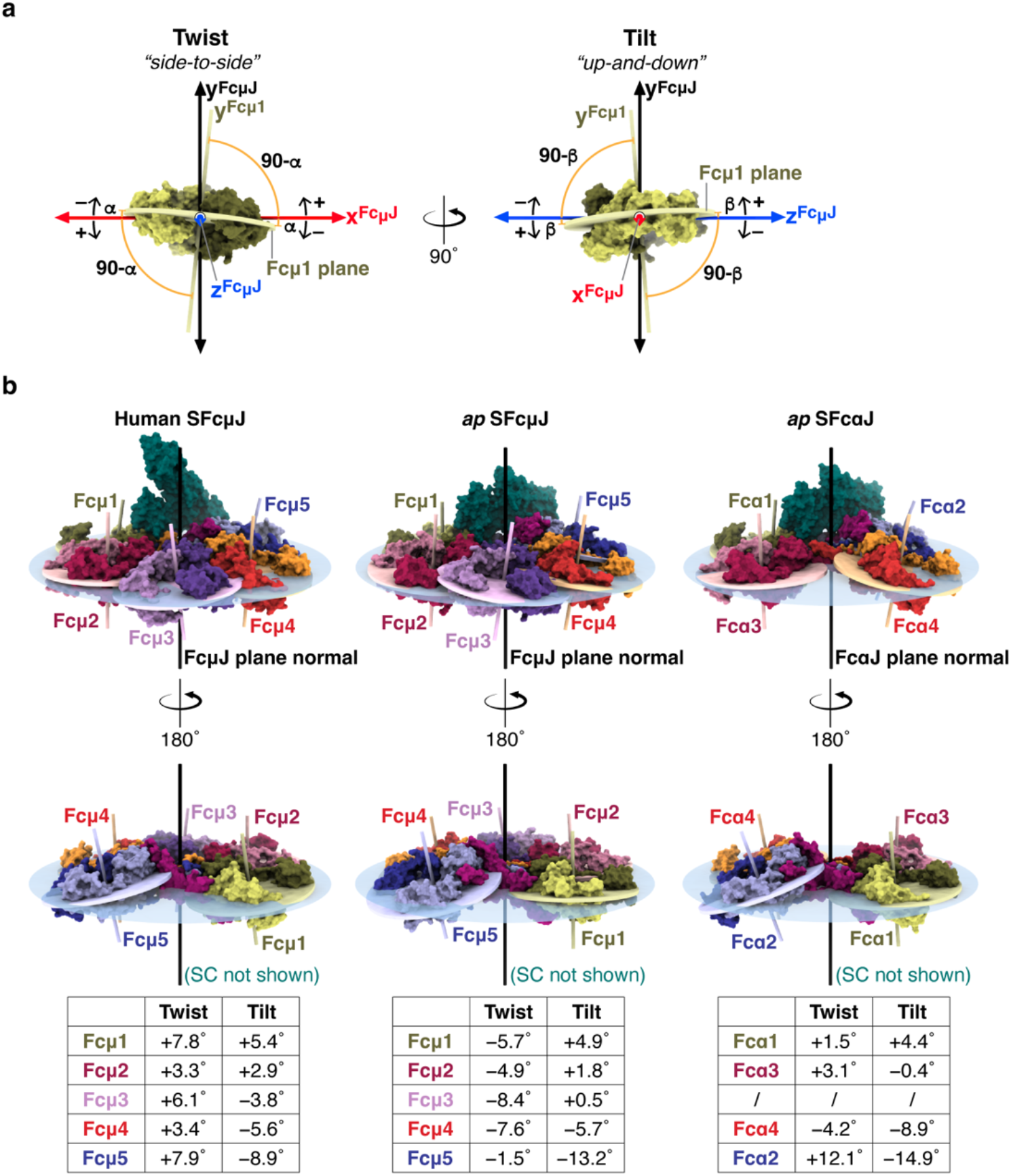
Twist and tilt measurements of human SFcμJ, *ap* SFcμJ, and *ap* SFcαJ. (**a**) Schematic representation of twist and tilt. Twist: angular deviation (α) of the subunit plane from the FcμJ plane, measured as the angle between the subunit plane (colored ellipses) and the fixed x^FcμJ^ axis (red line). Tilt: angular deviation (β) of the subunit plane from the FcμJ plane, measured as the angle between the subunit plane (colored ellipses) and the fixed z^FcμJ^ axis (blue line). (**b**) Molecular surface representations of human SFcμJ (PDB 6KXS; left), *ap* SFcμJ (center), and *ap* SFcαJ (PDB 9EC6; right) shown from two views, with (top) and without (bottom) SC. The FcμJ/FcαJ central planes (light blue, semi-transparent ellipses), FcμJ/FcαJ central plane normals (black), Fcμ/Fcα subunit planes (colored ellipses), and Fcμ/Fcα subunit plane normals (colored lines) are also shown. Twist and tilt values for each subunit are provided in tables below.

**Supplementary Figure 5.**
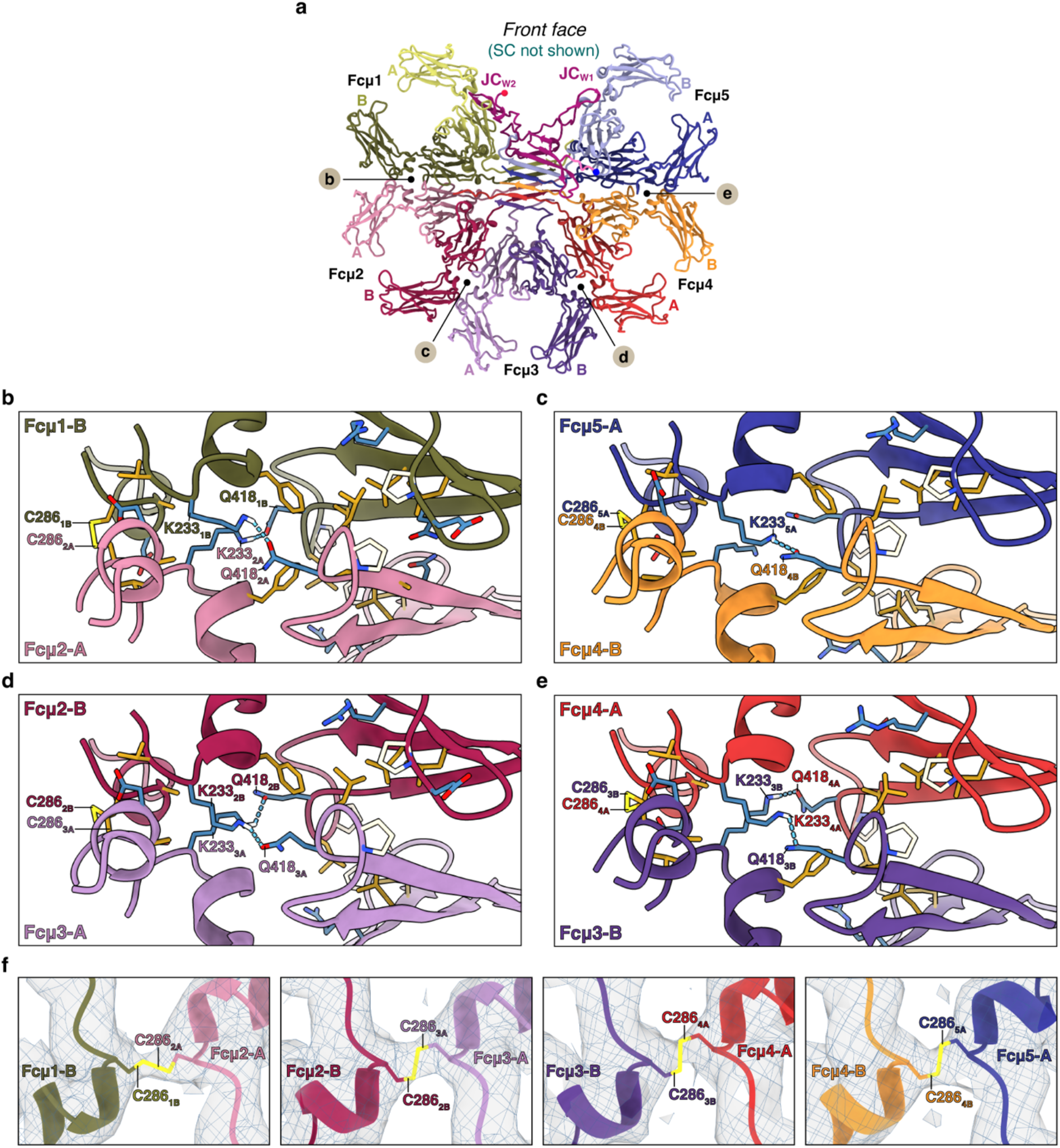
Fc-Fc interfaces of *ap* SFcμJ. (**a**) Cartoon representation of *ap* SFcμJ front face views; colored same as in (Fig. 1) with SC removed. JC N– and C-termini are indicated by blue and red spheres, respectively; JC_W1_ and JC_W2_ are labeled. Letters in circles indicate the regions enlarged in corresponding figure panels. (**b**-**e**) Fcμ1-Fcμ2 (b), Fcμ2-Fcμ3 (c), Fcμ3-Fcμ4 (d), and Fcμ4-Fcμ5 (e) interfaces shown as cartoon with residues involved in contacts shown as sticks and colored either gold (hydrophobic), cream (neutral), or blue (hydrophilic). Residues participating in interchain disulfide or hydrogen bond are labeled. (**f**) Cartoon representation surrounding Cys286 inter-chain disulfides observed at each Fc-Fc interface with Cys shown as sticks. Cryo-EM map density is shown using blue mesh and a light grey transparent surface, contoured to a threshold of 0.14 and carved at 2 Å.

**Supplementary Figure 6.**
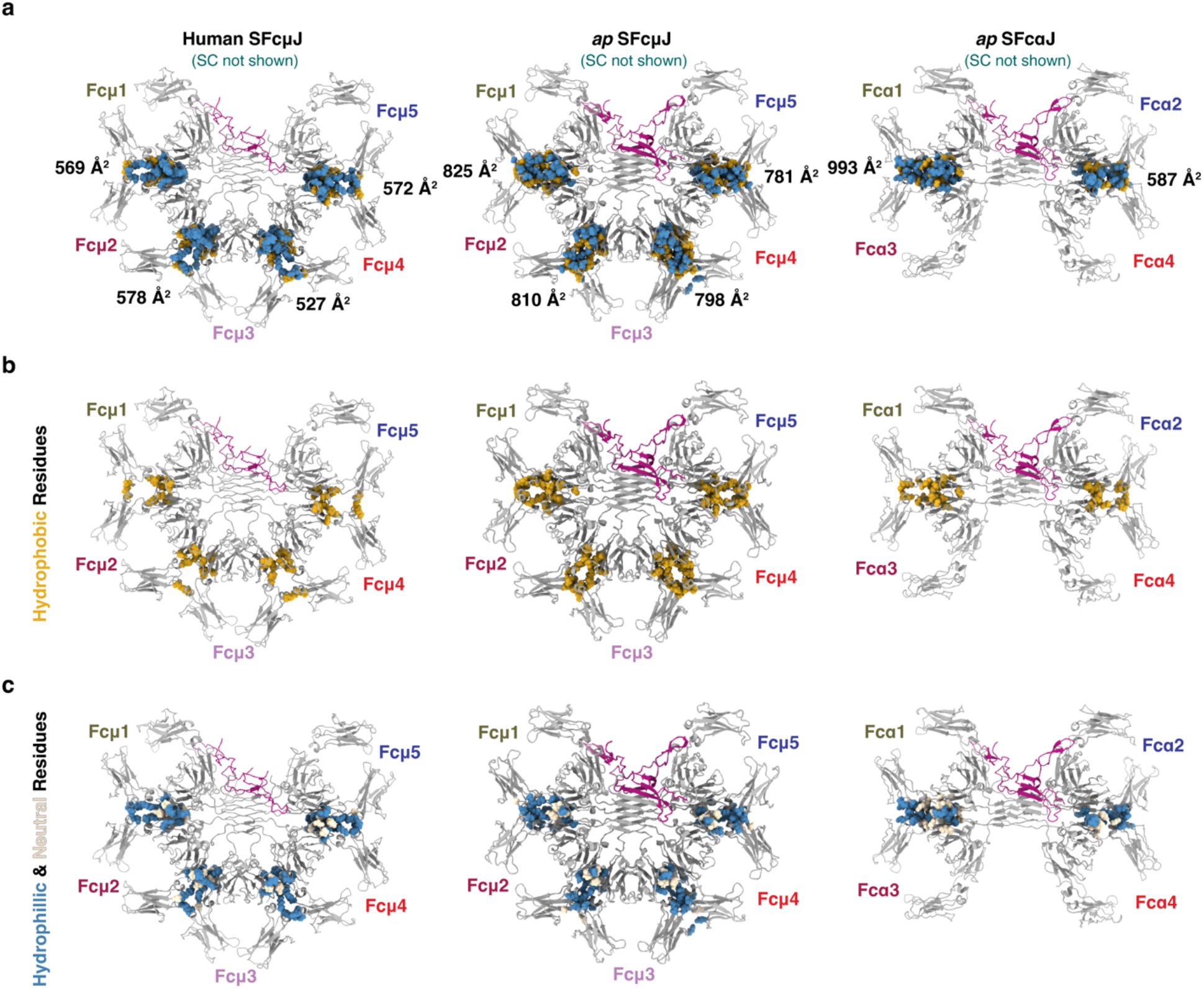
Contacting residues at Fc-Fc interfaces in human SFcμJ, *ap* SFcμJ, and *ap* SFcαJ. (**a**) Cartoon representations of human SFcμJ (PDB 6KXS; left), *ap* SFcμJ (center), and *ap* SFcαJ (PDB 9EC6; right) without SC; HCs are colored grey and JC colored magenta. Residues involved in Fc-Fc contacts shown as spheres and colored either gold (hydrophobic), cream (neutral), or blue (hydrophilic). Fc subunits are labeled and buried surface areas for Fc-Fc interfaces are shown in black text. (**b**) Same as (a), with only hydrophobic residues at Fc-Fc interfaces shown. (**c**) Same as (a), with only neutral and hydrophilic residues at Fc-Fc interfaces shown.

**Supplementary Figure 7.**
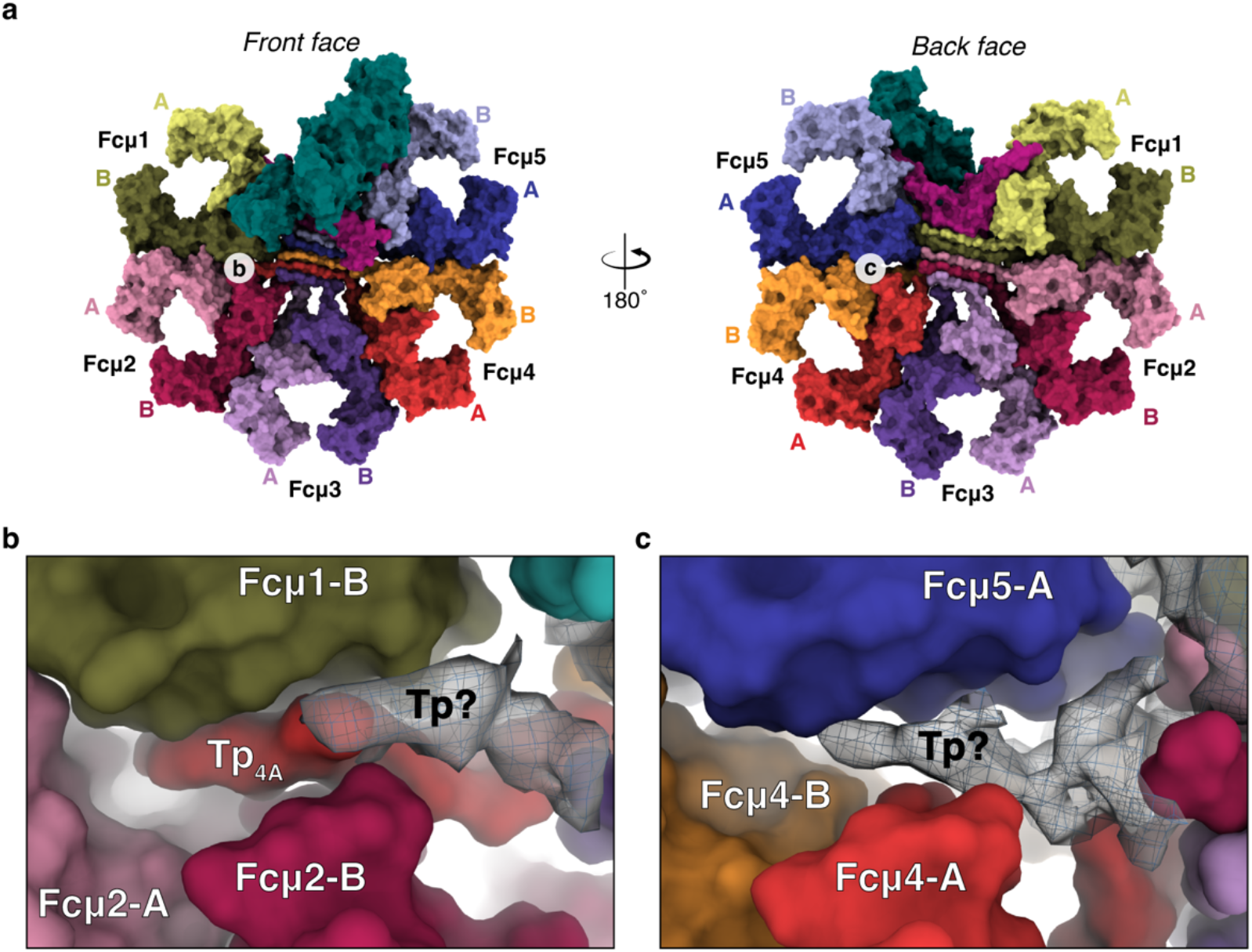
Putative Fcμ-Fcμ-Tp interfaces of *ap* SFcμJ. (**a**) Surface representation of *ap* SFcμJ front and back faces colored as in (Fig. 1). Fcμ subunits and chains are labeled. Circled letters indicate the corresponding figure panels that show enlarged views of the area. (**b**) Surface representation of Fcμ1-Fcμ2-Tp_4A_ interface. Unassigned cryo-EM map density is shown using blue mesh and a light grey transparent surface, contoured to a threshold of 0.14 and carved at 2 Å. (**c**) Surface representation of Fcμ4-Fcμ5 interface. Unassigned cryo-EM map density is shown using blue mesh and a light grey transparent surface, contoured to a threshold of 0.14 and carved at 2 Å.

**Supplementary Figure 8.**
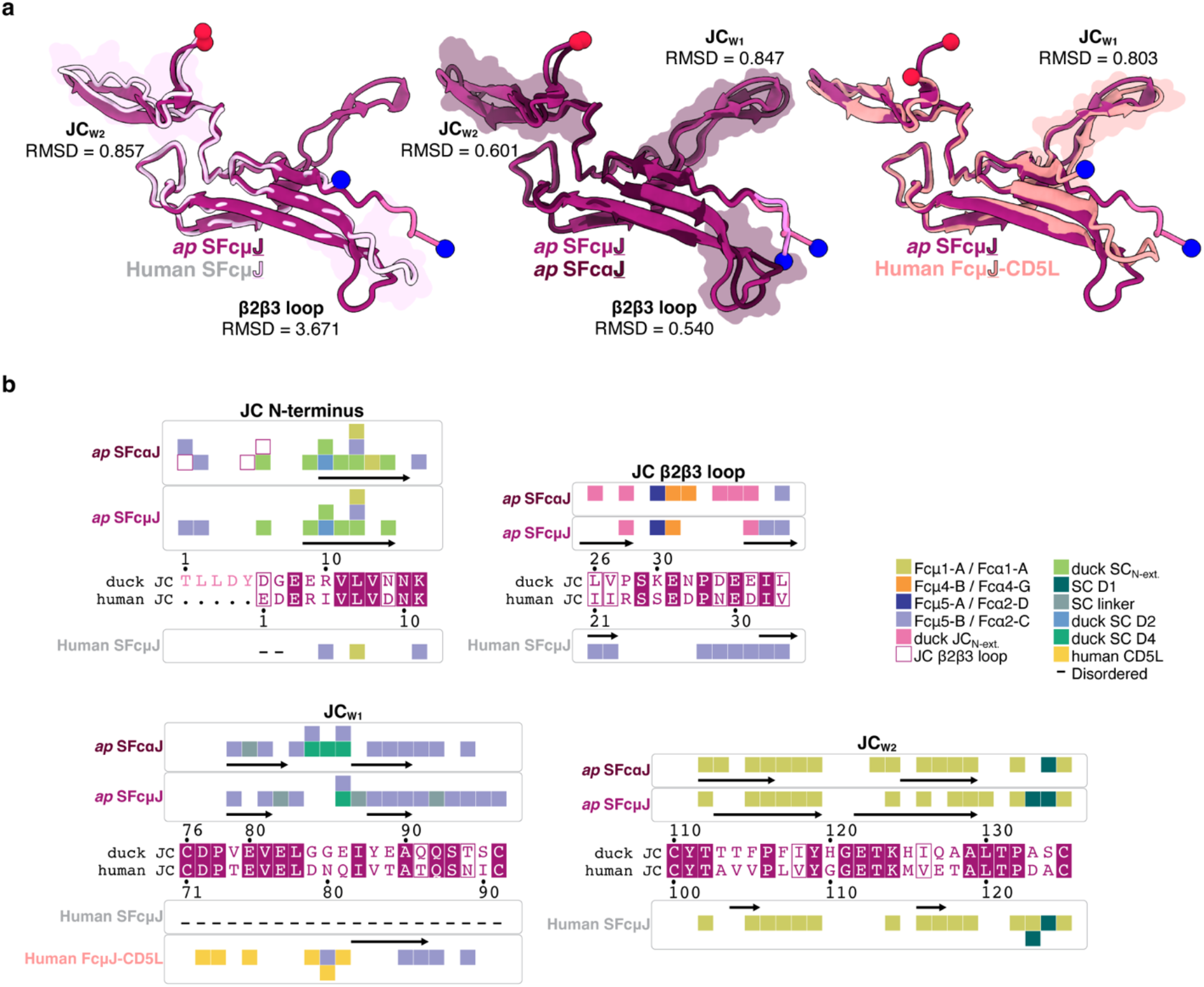
Structure-based comparisons of JC in human SFcμJ, *ap* SFcμJ, and *ap* SFcαJ. (**a**) Cartoon representations of JC from *hs* SFcμJ (PDB 6KXS; left), *ap* SFcαJ (PDB 9EC6; center), and human FcμJ-CD5L (PDB 8WYR; right) aligned to JC from *ap* SFcμJ. JC N– and C-termini are indicated by blue and red spheres, respectively. JC structural motifs are labeled with RMSD indicated and regions used to calculate values shown as flat surface representations. (**b**) Structure-based sequence alignments of duck and human JC motifs are shown with contacts for each structure in (a) indicated by colored squares according to key. Dashes signify disordered residues.

**Supplementary Figure 9.**
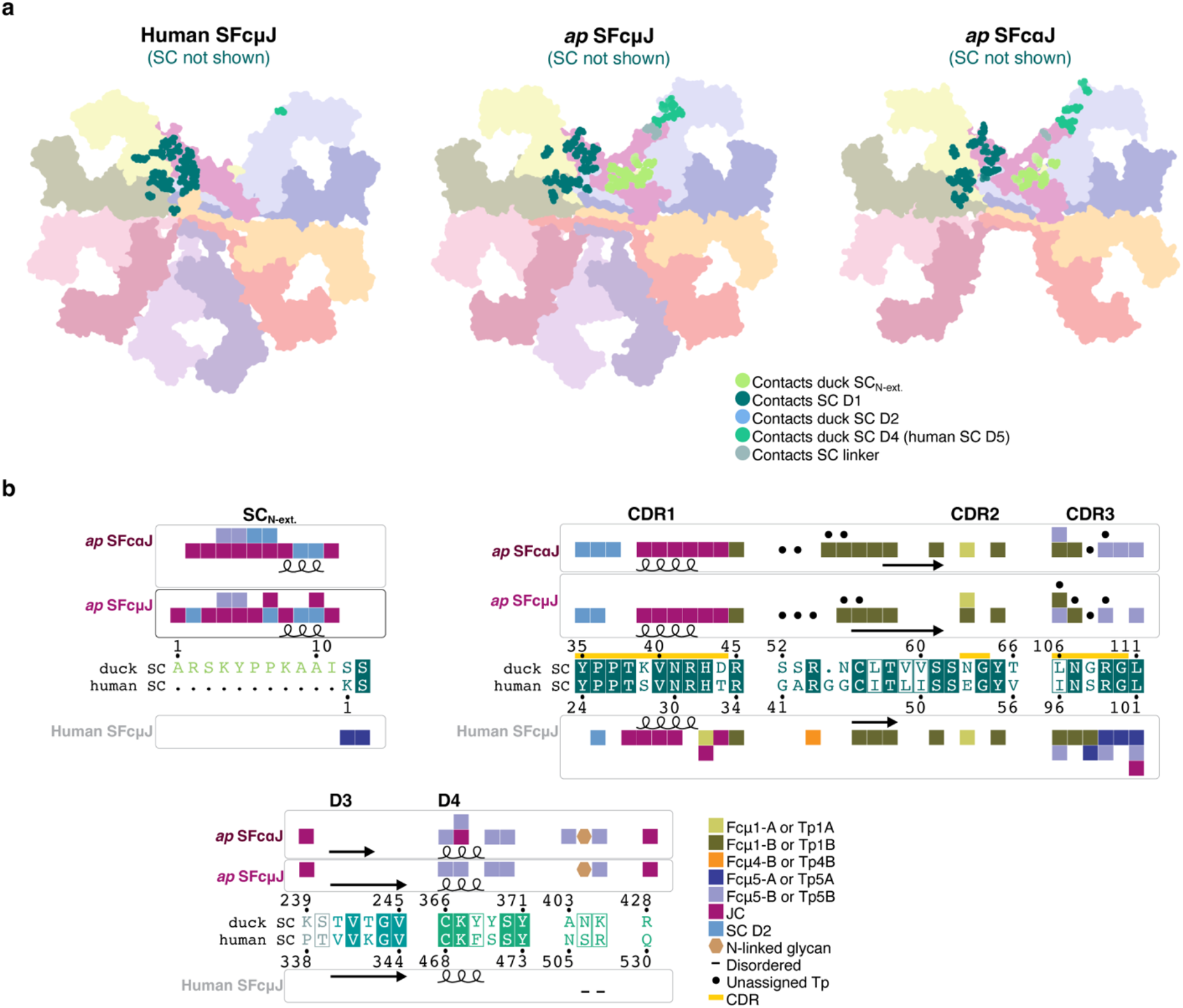
Comparison of SC binding in human SFcμJ, *ap* SFcμJ, and *ap* SFcαJ. (**a**) Flat surface representation of human SFcμJ (PDB 6KXS; top), *ap* SFcμJ (center), and *ap* SFcαJ (PDB 9EC6; bottom) with SC removed. HC and JC residues contacting SC are colored according to the key. (**b**) Structure-based sequence alignments of duck and human SC motifs are shown with colored squares to indicate contacts with HCs, JC, or other SC domains for each structure in (a) according to the key. Dashes signify disordered residues.

**Supplementary Figure 10.**
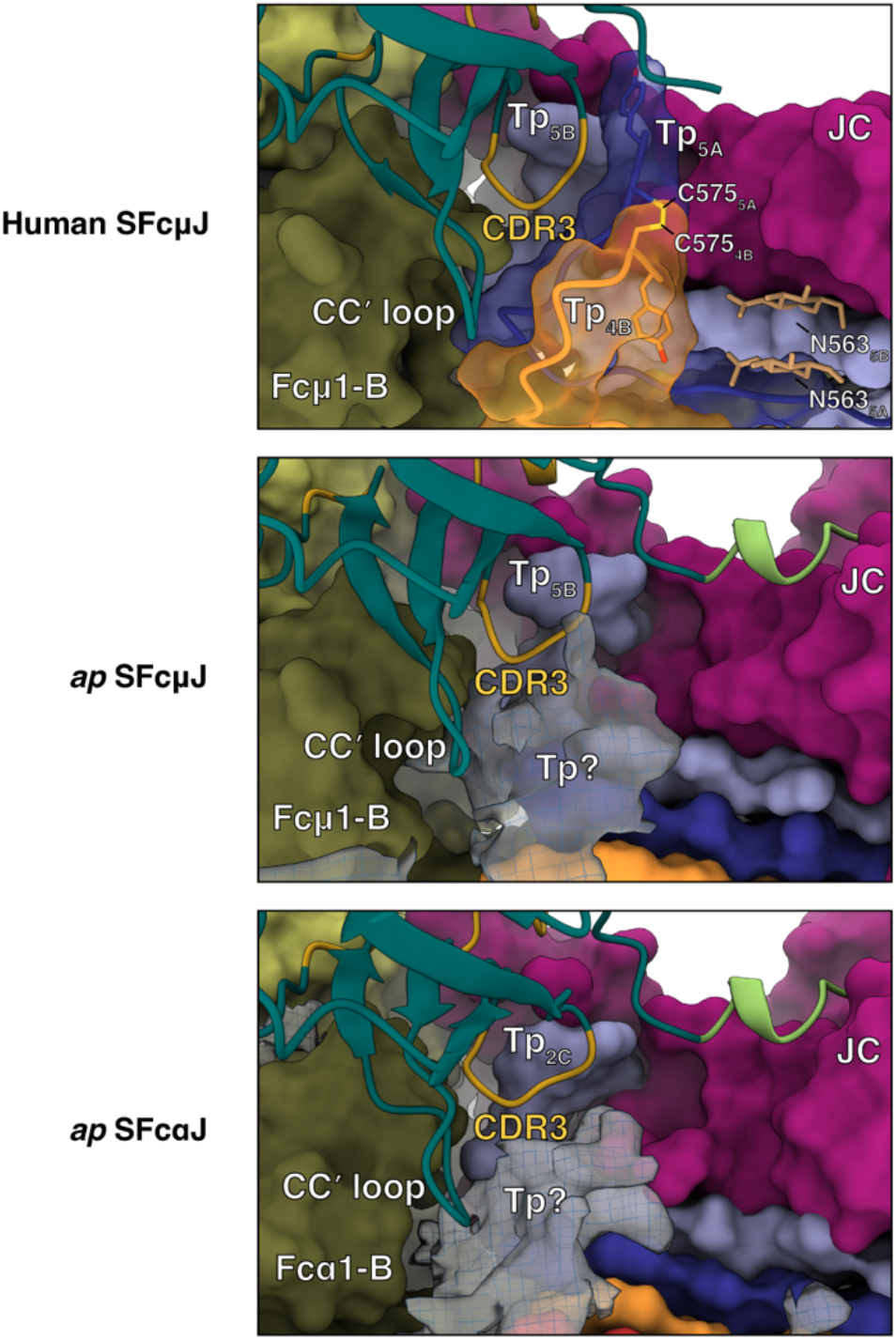
D1-Tp interfaces in human SFcμJ, *ap* SFcμJ, and *ap* SFcαJ. Surface representations of HCs and JC in human SFcμJ (PDB 6KXS; top), *ap* SFcμJ (center), and *ap* SFcαJ (PDB 9EC6; bottom) with SC D1 shown as a cartoon. In human SFcμJ, Tp_4B_ and Tp_5A_ surfaces are transparent and shown as cartoons with Cys575 and Tyr576 shown as sticks; glycans are also shown as sticks. Unassigned cryo-EM map density for *ap* SFcμJ and *ap* SFcαJ (EMD-47900) are shown using blue mesh and a light grey transparent surface, contoured to a threshold of 0.14 and carved at 2 Å.

**Supplementary Figure 11.**
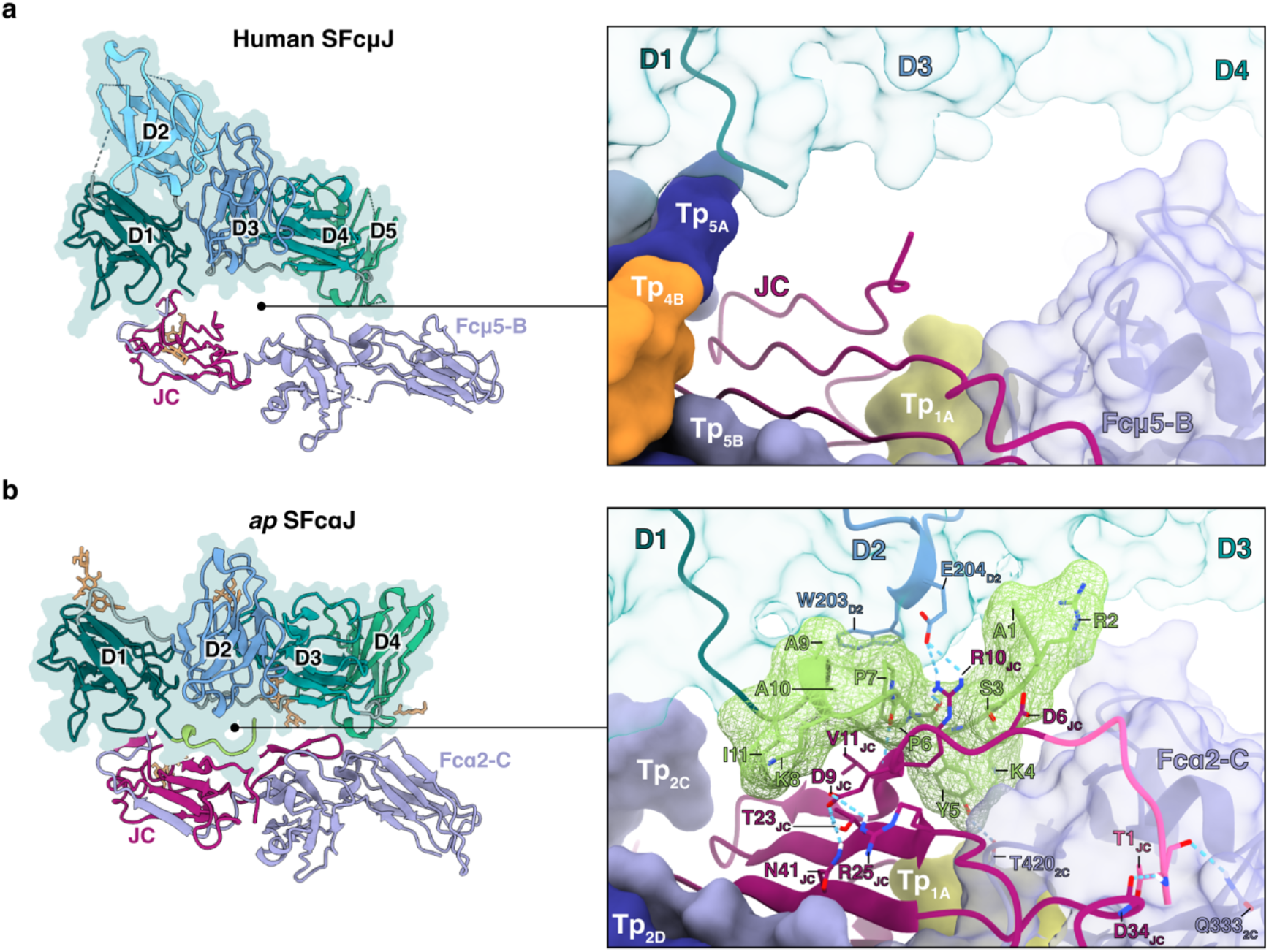
SC and JC N-termini in human SFcμJ and *ap* SFcαJ. (**a**) Left, cartoon representations of SC, Fcμ5-B, and JC only from human SFcμJ (PDB 6KXS) uniquely colored and labeled with glycans shown as sticks. Right, molecular surface representation of SC (semi-transparent), Fcμ5-B (semi-transparent), Tp_1A_, Tp_4B_, Tp_5A_, and Tp_5B_. JC and the SC N-terminus are shown as a cartoon. (**b**) Left, cartoon of SC, Fcα2-C, and JC only from *ap* SFcαJ (PDB 9EC6) uniquely colored and labeled with glycans shown as sticks. Right, molecular surface representation of SC (semi-transparent), SCN-ext. (mesh), Fcα2-C (semi-transparent), Tp_1A_, Tp_2C_, and Tp_2D_. SC_N-ext._, a D2 loop, and JC are shown as cartoons; the JC_N-ext._ is uniquely colored. Contacting residues are shown as sticks and labeled.

**Supplementary Figure 12.**
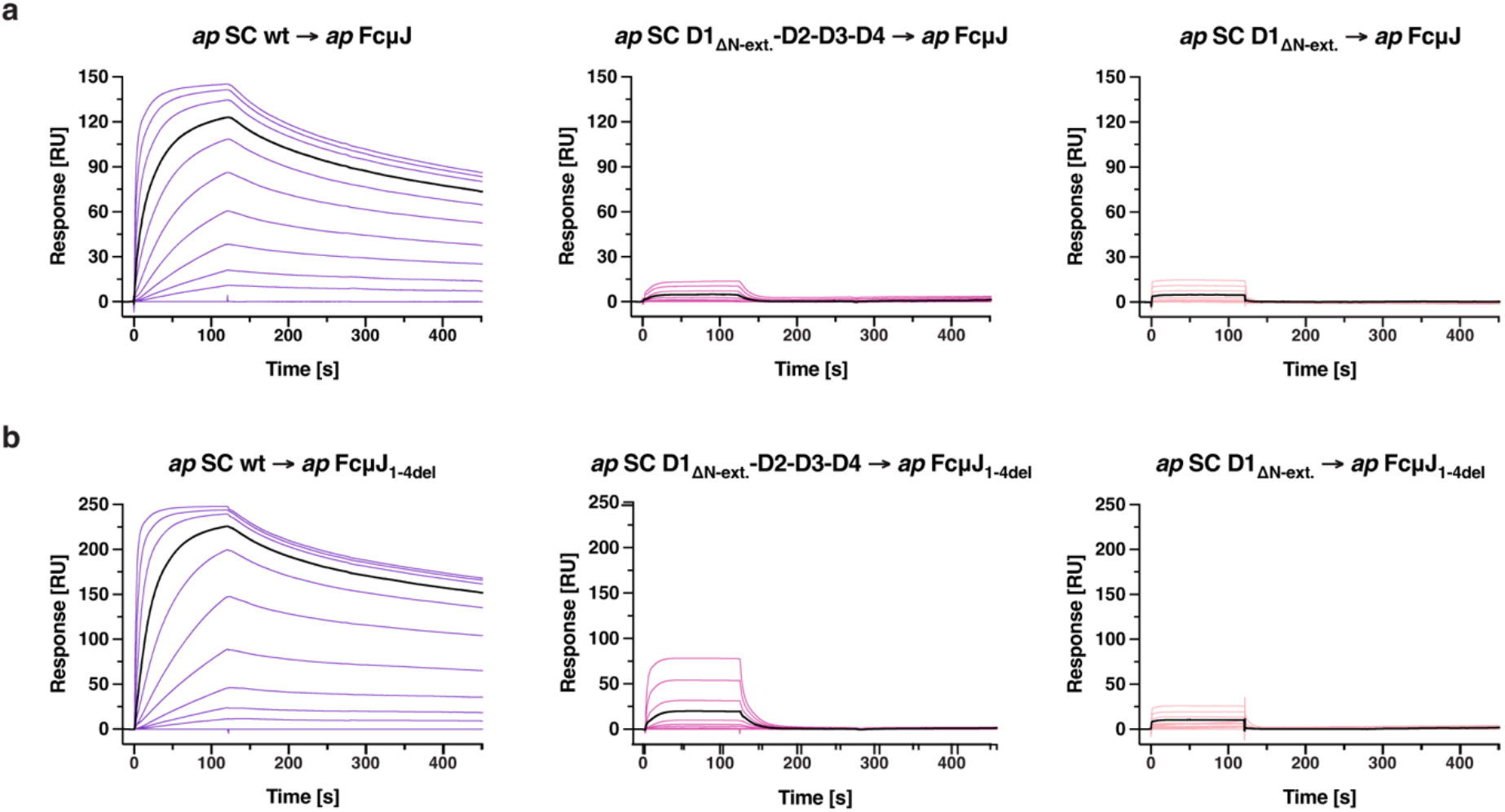
Duck SC variants binding to *ap* FcμJ. (**a**, **b**) SPR sensorgrams showing responses for entire two-fold dilution series of *ap* SC variants binding to *ap* FcμJ (a) and *ap* FcμJ_1-4del_ (b). The maximum concentration used was 512 nM for all analytes. Sensorgrams are colored to match corresponding curves in concentration-matched panels (Fig. 5), except for responses at 64 nM which are colored black. Sensorgrams shown are equivalent to those obtained from replicate experiments (not shown).

**Supplementary Figure 13.**
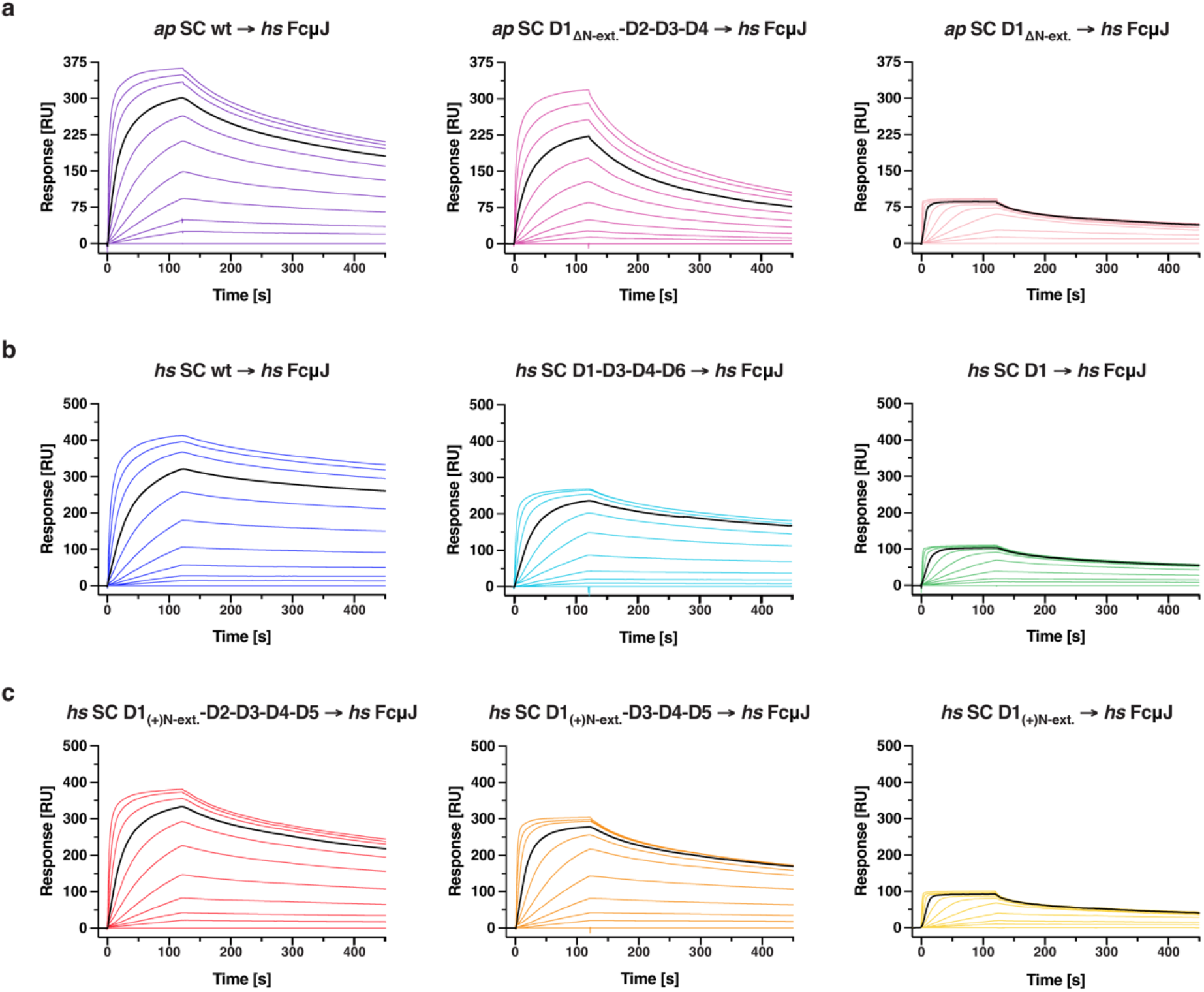
Duck and human SC variants binding to *hs* FcμJ. (**a**-**c**) SPR sensorgrams showing responses for entire two-fold dilution series of *ap* SC (a), *hs* SC (b), or chimeric *hs* SC (c) variants binding to *hs* FcμJ. The maximum concentration used was 512 nM for all analytes. Sensorgrams are colored to match corresponding curves in concentration-matched panels (Fig. 5), except for responses at 64 nM which are colored black. Sensorgrams shown are equivalent to those obtained from replicate experiments (not shown).

**Supplementary Figure 14.**
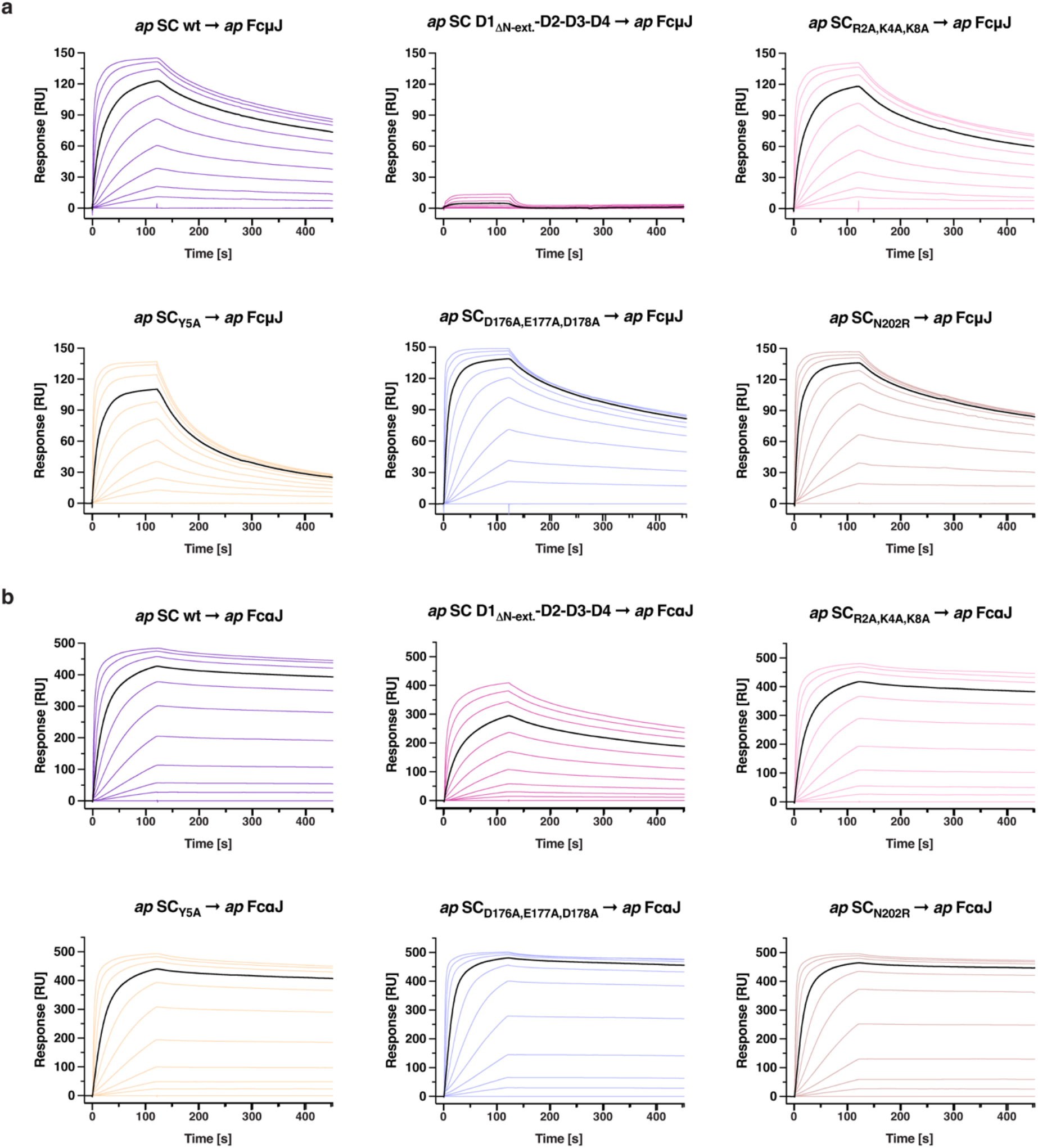
Duck SC variants binding to *ap* FcμJ and *ap* FcαJ. (**a**, **b**) SPR sensorgrams showing responses for entire two-fold dilution series of *ap* SC variants binding to *ap* FcμJ (a) and *ap* FcαJ (b). The maximum concentration used was 512 nM for all analytes. Sensorgrams are colored to match corresponding curves in concentration-matched panels (Fig. 5), except for responses at 64 nM which are colored black. Sensorgrams shown are equivalent to those obtained from replicate experiments (not shown).

**Supplementary Figure 15.**
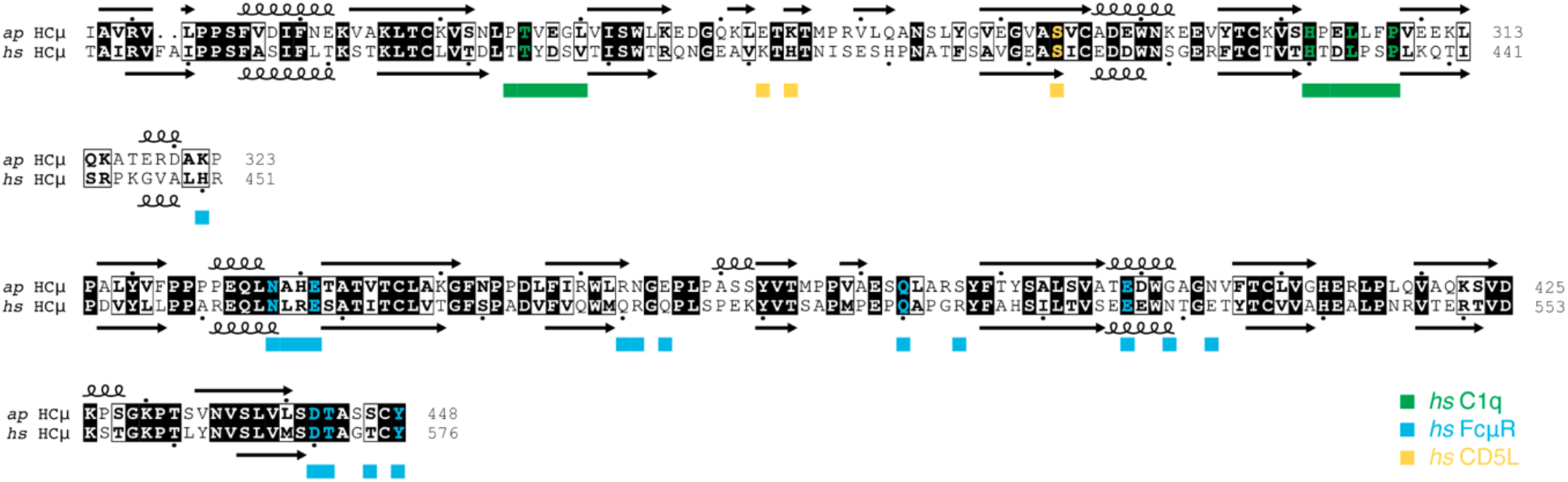
Conservation of C1q, FcμR, and CD5L binding sites in duck IgM HC. Sequence alignment of duck IgM heavy chain (*ap* HCμ) with human heavy chain IgM (*hs* HCμ). Human HCμ residues contacting C1q, FcμR, or CD5L are indicated by green, blue, or yellow squares, respectively. Conserved contacting residues in duck HCμ are bold and colored to indicate the corresponding receptor.

**Supplementary Table 1.** Cryo-EM data collection and model refinement statistics.

|  | *ap* SFcμJ<br>EMD-75637<br>PDB 11DN |
| --- | --- |
| **Data collection** |  |
| Voltage (kV) | 300 |
| Nominal magnification | 81,000x |
| Total dose (electrons/Å<sup>2</sup>) | 57.79 |
| Total exposure time (s) | 3.21 |
| Raw movie pixel size (Å/pixel) | 0.527 |
| Defocus range (μm) | 0.6 – 2.8 |
| Frames per movie | 40 |
| Automation software, 0°/30° tilt | EPU/Leginon |
| **Data processing** |  |
| Movies collected, 0°/30° tilt | 3,631 / 3,632 |
| Initial particles | 12.1M |
| Final particles | 431,415 |
| Map resolution (Å) | 3.37 |
| FSC threshold | 0.143 |
| **Model fit and validation** |  |
| **Model vs. data (CC)** |  |
| Mask | 0.82 |
| Box | 0.88 |
| Peaks | 0.79 |
| Volume | 0.81 |
| **Model composition** |  |
| Total atoms | 44,291 |
| Non-hydrogen atoms | 22,213 |
| Protein residues | 2,850 |
| Ligands | BMA: 1; NAG:8 |
| **Bond RMSD** |  |
| Lengths (Å) | 0.003 |
| Angles (°) | 0.555 |
| MolProbity score | 1.43 |
| Clash score | 4.74 |
| **Ramachandran plot (%)** |  |
| Outliers | 0.00 |
| Allowed | 3.08 |
| Favored | 96.92 |
| Rotamer outliers (%) | 0.00 |
| Cβ outliers (%) | 0.00 |
| Cis proline | 0.0 |
| Twisted proline | 0.0 |
| CaBLAM outliers (%) | 2.14 |

**Supplementary Table 2.**
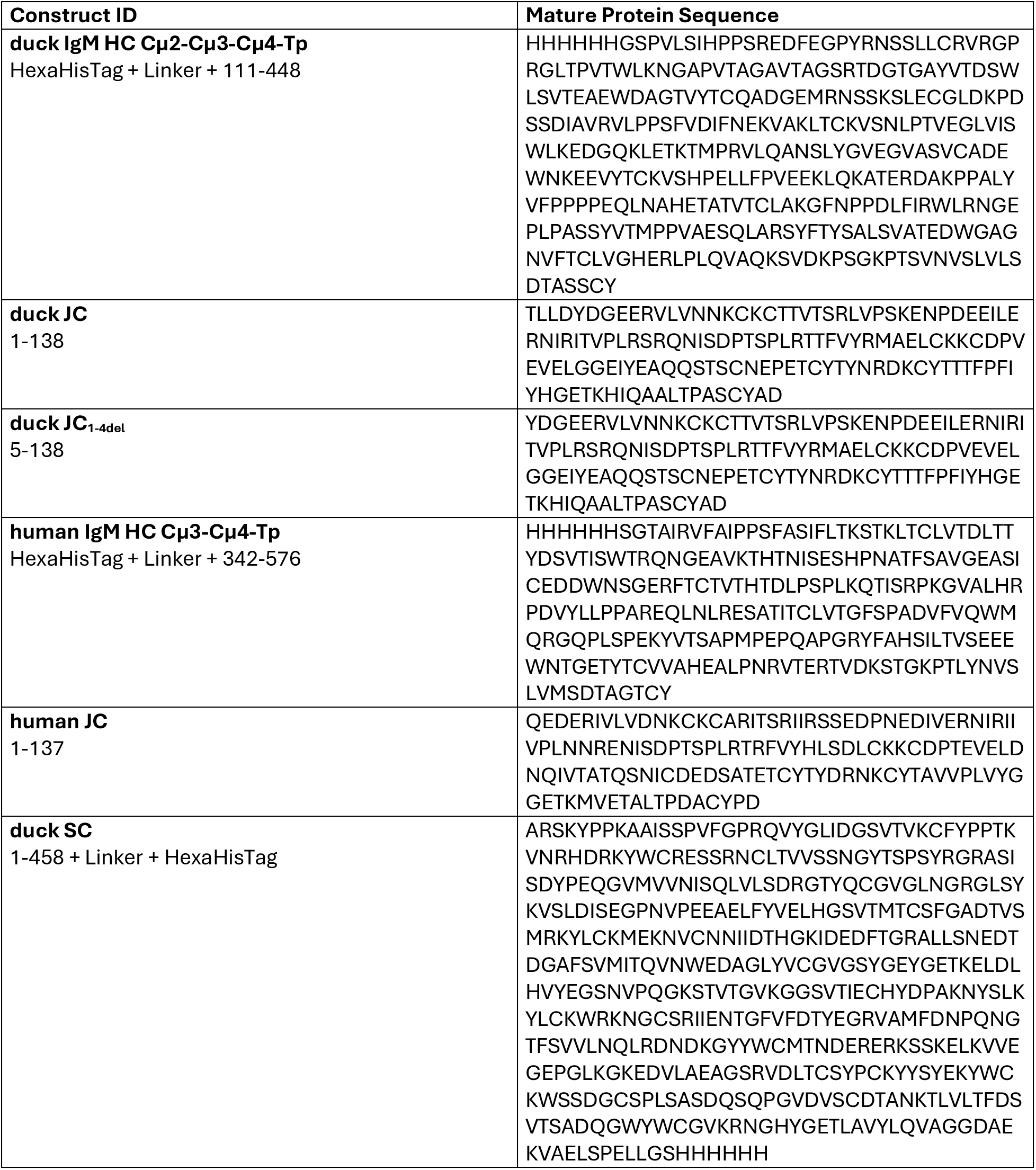
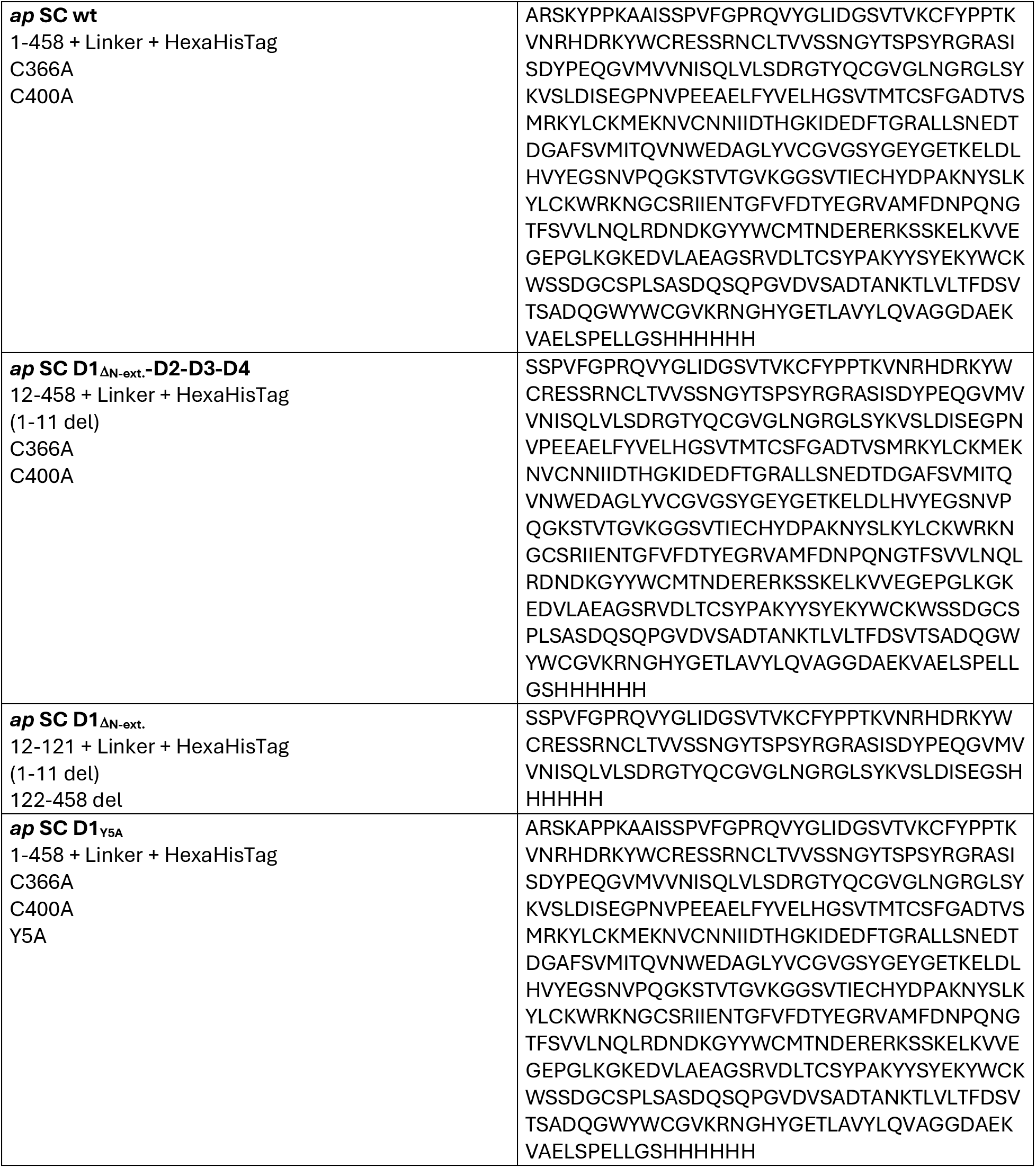
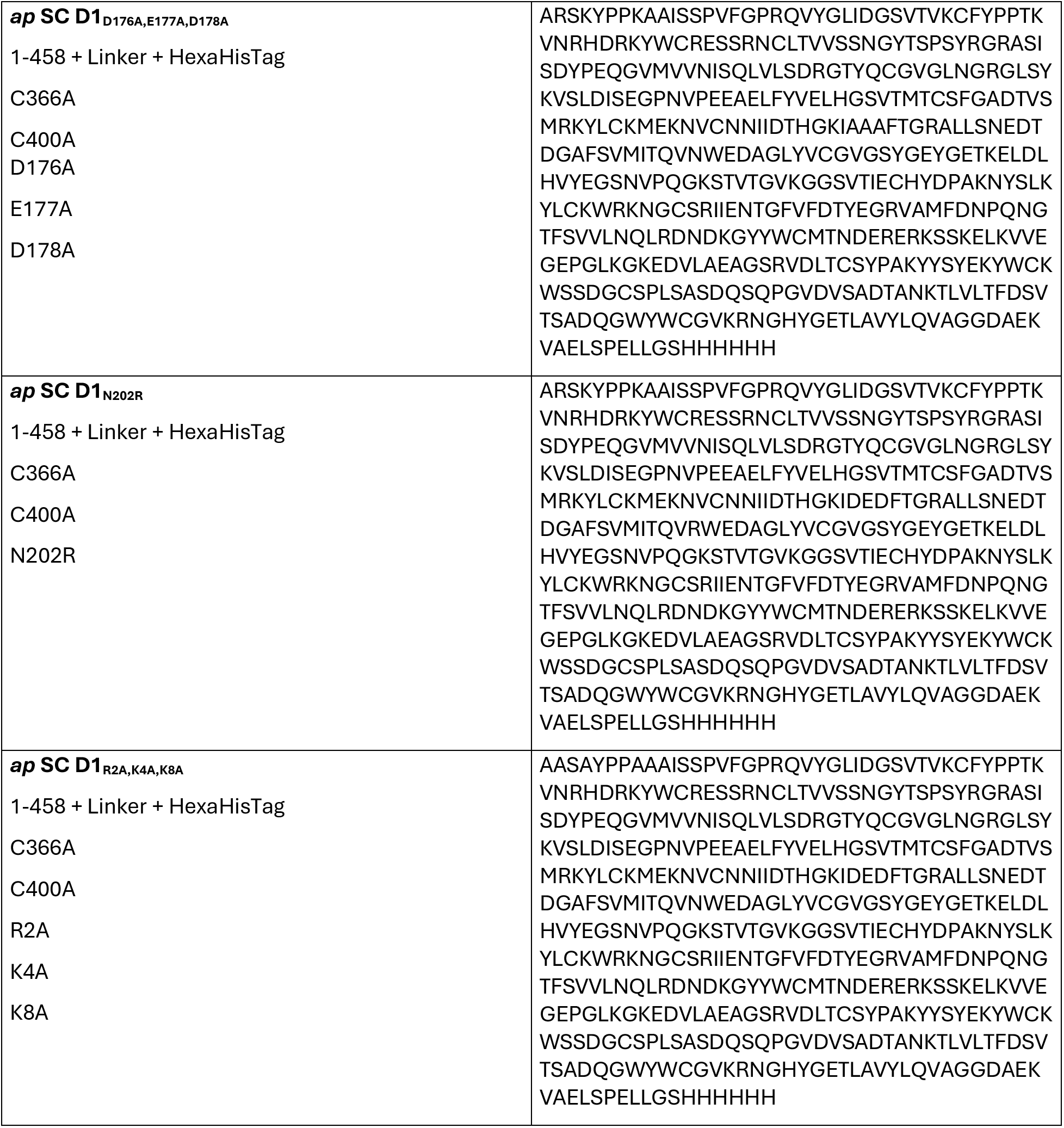
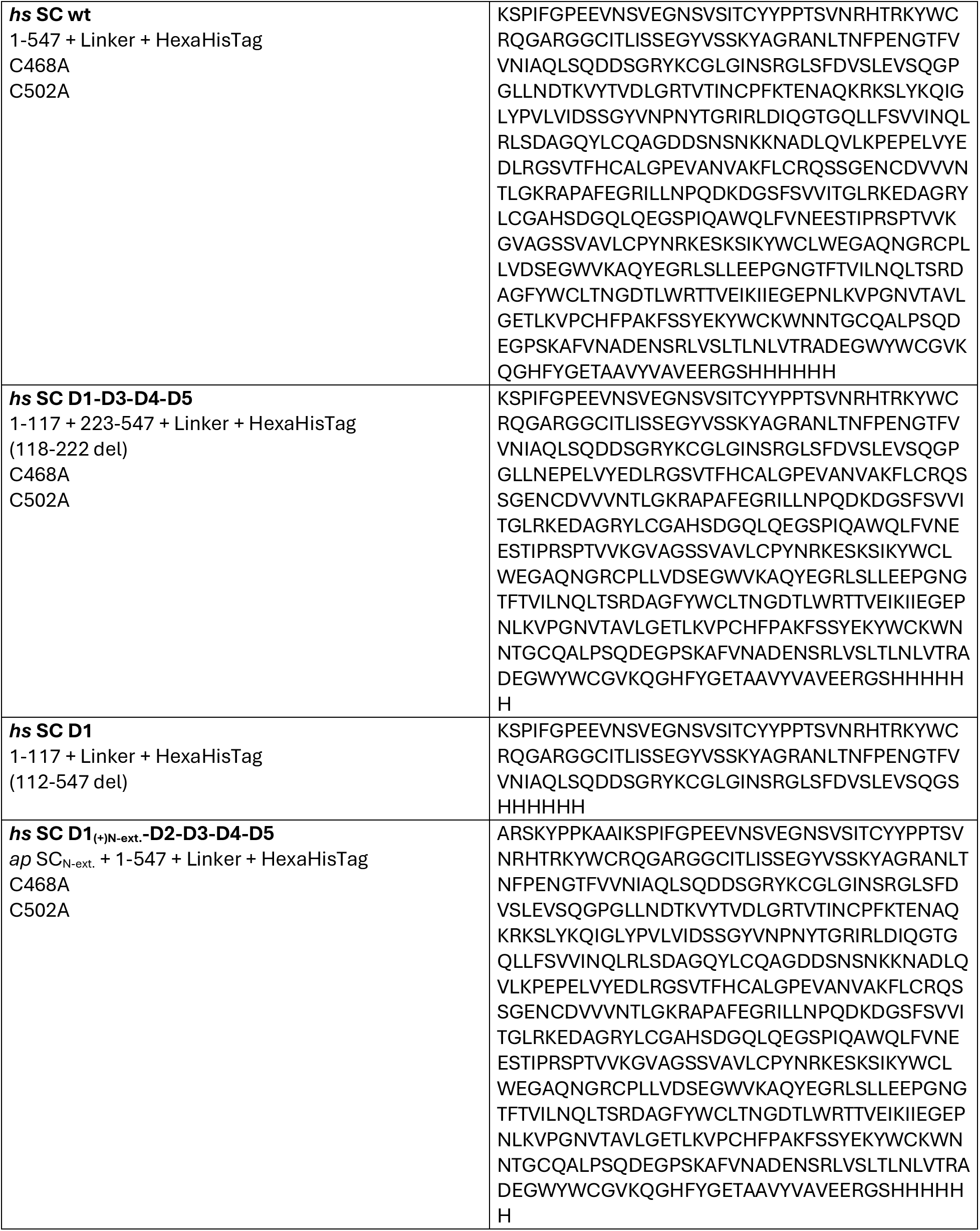
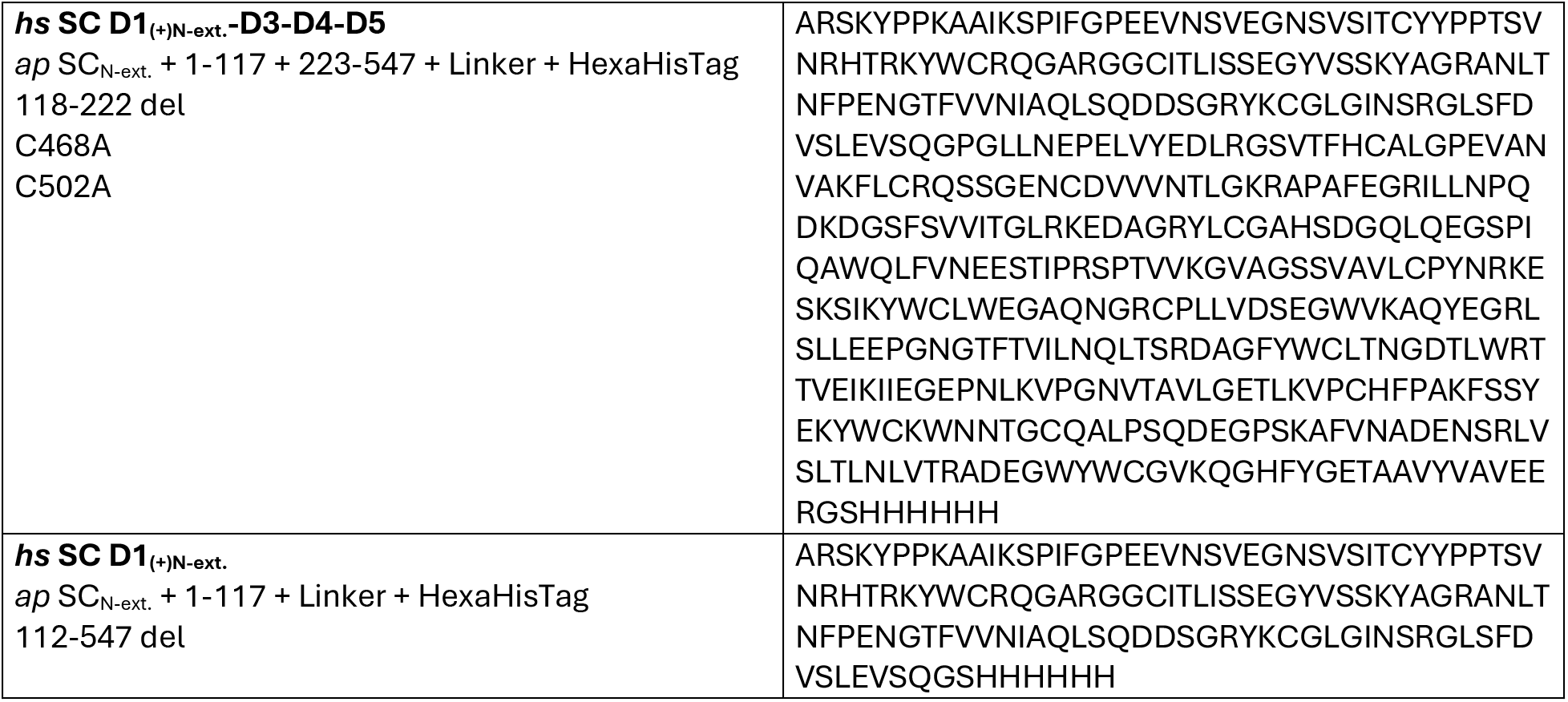
Expression constructs used in this study.

